# Intracellular trafficking of an Arabinogalactan protein SLEEPING BEAUTY influences cell wall integrity and apical tip growth in *Physcomitrium patens*

**DOI:** 10.1101/2025.07.08.663806

**Authors:** Chao-Yuan Yu, Luis Alonso Baez, Manju Maharjan, Chao-Hsi Liu, Thorsten Hamann, Ooi-Kock Teh

## Abstract

Polarized tip growth is a key morphological adaption that facilitated land plant terrestrialization, and arabinogalactan proteins (AGPs) are central regulators of this process. However, how glycosylation modulates AGP function during tip growth remains poorly understood. Here, we show that the AGP SLEEPING BEAUTY (SB) is required for cell wall integrity during protonemal tip growth in the moss *Physcomitrium patens*, a function that is conserved in the distantly related vascular plant *Arabidopsis thaliana*. Loss of SB function disrupts cellulose microfibril organization and compromises cell wall integrity, leading to altered tip growth dynamics and geometry. Using a hypoglycosylated SB variant, we demonstrate that glycosylation of Pro55, Pro92, and Pro94 within a highly intrinsically disordered region is required for proper SB secretion and vacuolar trafficking; loss of these modifications results in pronounced inhibition of tip growth. We propose that glycosylation of the intrinsically disordered domain in AGP fine-tunes SB abundance at the cell surface, thereby facilitating cellulose microfibril biosynthesis and/or assembly to regulate robust polarized tip growth.

## Introduction

All land plants share several morphological innovations traceable to the last common ancestor of embryophytes, among which apical tip growth is a defining feature. Thought to have arisen during the aquatic-to-terrestrial transition, tip growth occurs in bryophyte rhizoids as well as in pollen tubes and root hairs of vascular plants. Unlike diffusive growth, where expansion occurs over the entire cell surface, tip growth is characterized by highly localized and rapid expansion at a single, restricted site, producing elongated filamentous structures. This growth mode is widely considered a key evolutionary adaptation that enhanced substrate anchorage and nutrient acquisition in early land plants.

In yeast and anumals, specification of the growing tip requires precise regulation by members of the highly conserved Cdc42/Rho/Rac small GTPase family(Park and Bi, 2007). Although this family is absent in plants, their roles are fulfilled by a plant-specific group of GTPases known as RHO OF PLANTS (ROP). In *Arabidopsis* and *Physcomitrium patens (P. patens)*, ROPs accumulate polarly at incipient root hair initiation sites and protonemal branching points, respectively (Molendijk et al., 2001; Yi and Goshima, 2020). Their spatially restricted localization triggers signaling cascades involving Ca²⁺ dynamics, cytoskeletal remodeling, vesicle trafficking, and exocytosis (Gu et al., 2005; Lee et al., 2008; Zhou et al., 2015). Exocytosis is particularly important for delivering cell wall materials such as highly methyl-esterified pectins. To enable rapid cellular expansion, the wall of tip-growing cells must balance elasticity—permitting rapid extension—with sufficient stiffness to withstand internal turgor pressure (reviewed in (Dehors et al., 2019)). This is achieved by secretion of newly synthesized pectin followed by localized de-esterification by pectin methylesterases, producing calcium–crosslinked homogalacturonan (“egg-box” structures). In rapidly growing pollen tubes, lowly-esterified “hard” pectin is enriched in the shank, whereas highly esterified “soft” pectin accumulates at the apex (Luo et al., 2017). Comparable tip-focused distribution patterns have not been reported for cellulose or hemicellulose, although inhibiting cellulose synthesis with 2,6-dichlorobenzonitrile (DCB) causes severe morphological defects and ultimately tip rupture (Anderson et al., 2002; Hao et al., 2013).

Beyond polysaccharides, cell wall structural proteins such as arabinogalactan proteins (AGPs) also play important, though elusive, roles in tip growth. AGPs are ubiquitous and conserved across land plants and contribute to a wide range of physiological processes. Their importance in tip growth is supported by their apical enrichment, inhibition of tip growth by the AGP-binding β-Yariv reagent (Lee et al., 2005), and genetic evidence from AGP mutants in *Arabidopsis* and *P. patens* (Lee et al., 2005; Borassi et al., 2020). AGP function is thought to be conferred largely by its complex glycan moieties, whose post-tanslational modifiction begins in endoplasmic reticulum (ER). Following co-translational insertion into the ER via an N-terminal localised signal peptide, proline residues in the AGP peptide backbone are hydroxylated by ER-localized prolyl hydroxylases (Yuasa et al., 2005; Velasquez et al., 2011). In many AGPs, the polypeptide is further modified by the addition of a glycosylphosphatidylinositol (GPI) anchor; notably 55 of the 85 annotated Arabidopsis AGPs contain a predicted GPI-attachment signal (Showalter et al., 2010). Subsequent glycosylation occurs in the Golgi apparatus, where a suite of arabinosyltransferases catalyze Hyp-arabinosylation of the peptide backbone, followed by the stepwise elongation and diversification of glycan side chains (Parsons et al., 2019; Silva et al., 2020). A longstanding question in glycobiology is how AGP glycosylation influences their function. Recent work demonstrates that β-linked glucuronic acid residues can bind calcium in a pH-dependent manner, enabling AGPs to act as extracellular calcium capacitors that facilitate calcium wave propagation(Lamport and Varnai, 2013; Lamport et al., 2014; Lopez-Hernandez et al., 2020). Emerging evidence also indicates that AGP glycosylation is required for their intracellular trafficking and secretion(Xue et al., 2017; Zhang et al., 2019).

We previously showed that two AGPs in *P. patens*, SLEEPING BEAUTY (SB) and its homolog SB-LIKE (SBL), function redundantly to modulate cell wall properties and thereby influence the timing of gametophore formation (Teh et al., 2022). Unlike canonical AGPs, SB is anchored to the plasma membrane using a transmembrane domain rather than a GPI lipid anchor (Teh et al., 2022). Further experiments identified AUXIN RESPONSE FACTOR C (ARFC) as a genetic interactor acting downstream of SB. However, the role of SB and SBL in cell wall integrity and remodelling remains unresolved, as the *sb sbl* double knockout displays no overt phenotype, whereas SB overexpression causes severe, dosage-dependent growth defects (Teh et al., 2022). We hypothesized that glycosylation is important for SB function. With this in mind, we identified key glycosylation sites and analyzed how impaired glycosylation affects SB intracellular trafficking. Our findings establish a mechanistic link between glycosylation-dependent AGP trafficking and cell wall integrity in the control of tip growth, providing a concrete example of how hydroxyproline (Hyp)-arabinoxylation contributes to AGP functionality.

## Results

### SB is glycosylated at P55, P92, and P94

Classical AGPs associate with the plasma membrane via glycosylphosphatidylinositol (GPI) anchors. In contrast, SB adopts a type-I membrane topology and contains a transmembrane domain. SB is also an intrinsically disordered protein as predicted by flDPnn, a computational tool that identifies intrinsic disorder in proteins (Hu et al., 2021). Its apoplastic-facing ectodomain is strongly enriched in proline residues, which constitute ∼23% of its amino acid composition. These prolines are likely hydroxylated to hydroxyprolines (Hyp) by prolyl 4-hydroxylases and subsequently serve as glycosylation sites for galactose addition by Hyp-O-galactosyltransferases, followed by elaboration of arabinan and arabinogalactan side chains(Petersen et al., 2021). Glycosylation is a major post-translational modification that substantially increases the molecular weight of AGPs. Consistent with this, fluorescent protein translation fusions such as SB-Dendra (Teh et al., 2022) and SB-mNeonGreen (SB-mNG, Fig. S1A) migrated as broad, high-molecular-weight smears of ∼90 kDa, double their predicted mass of 40 kDa. Such mobility shift could be caused by *N*- or *O*-linked glycosylation.To determine whether *N*-linked glycosylation contributed to this shift, we first looked for canonical *N*-glycosylation motifs in the SB ectodomain. SB lacks the Asn-X-Ser/Thr consensus sequence required for *N*-glycosylation, and its mobility was unaffected by treatment with *N*-glycosidases EndoH and PNGaseF (Fig.S1B). We therefore focused on identifying possible *O*-linked glycosylation sites in SB.

We previously demonstrated that mutating five of the 22 proline residues in SB did not altering the apparent molecular weight (Teh et al., 2022), suggesting that these proline residues are not Hyp-*O*-glycosylation sites. To identify bona fide Hyp-*O*-glycosylation sites in SB, we predicted potential hydroxylation sites using the [AVSG]-Pro-[AVST]-[AVPSTC]-[APS] motif proposed by Shimizu et al(Shimizu et al., 2005). Five proline residues within the SB ectodomain- P55, P92, P94, P96, P98- match this motif (Fig. 1A). Notably, P55, P92 and P94 correspond to regions with either the highest (P92, P94) or near-highest (P55) intrinsic disorder propensity (Fig. 1B), and subsituting these residues with alanine is predicted to decrease the disorder propensity (Fig. 1B). We therefore subsituted these proline residues with the nonpolar alanine and generated a series of mNG translational fusion reporters: SB^P55A^-mNG, SB^P55,92A^-mNG and SB^P55,92,94A^-mNG, alongside the WT reference SB-mNG. Because native promoter-driven SB is barely detectable and inducible SB overexpression caused severe growth defects (Teh et al., 2022), all constructs were expressed from a UBIQUITIN promoter in the WT background to produce semi-strong overexpressor lines. Next, we examined the molecular weight of these SB variants. All three variants displayed reduced molecular weight compared with SB-mNG (Fig. 1C); however, none migrated as low as the predicted size of fully unglycosylated SB-mNG (∼40kDa). To verify the migration behavior of unglycosylated SB, we expressed a His-tagged SB protein in glyscosylation-defective *E. coli*. SB-His migrated at its predicted molecular weight of19.2 kD (Fig. 1D). These results indicate that additional glycosylation sites remain in SB and that SB^P55,92,94A^-mNG represents hypoglycosylated-but not completely deglycosylated-variant.

**Figure 1.**
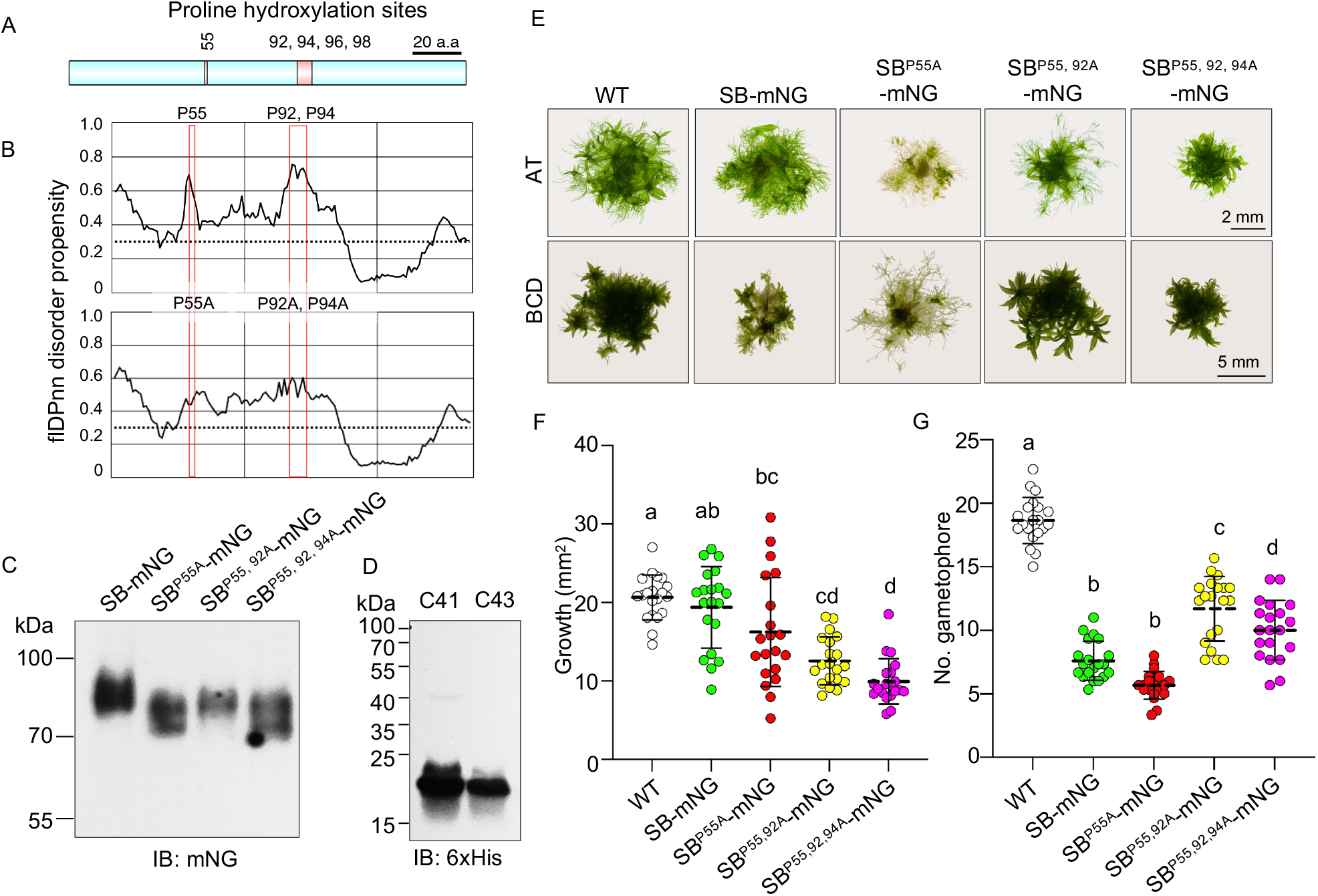
Generation and characterization of SB-mNG and hypoglycosylated SB reporter lines. (A) Schematic representation of SB showing putative proline hydroxylation sites (numbers above the cartoon). (B) Prediction of disordered propensities plots for SB (top) and SB^P55,92,94A^ (bottom). Red boxes indicate the mutated proline sites. Substitution of P55, P92 and P94 is predicted to reduce the protein disorder propensity. (C) Immunoblot analysis of mNG-immunoprecipitated proteins from SB-mNG and SB glycan variants reporter lines. Hypoglyscosylated SB variants exhibit reduced apparent molecular weight. (D) Immunoblot of recombinant His-tagged SB proteins expressed in *E.coli* C41 and C43 strains. Deglycosylated SB migrates at the predicted molecular weight. (E) Representative images of 21-day-old colonies of WT, SB-mNG and SB glycan variant reporter lines grown on AT (top) or BCD (bottom) medium. (F) Scatter dot plot showing colony area size after 21 days of growth on AT medium. WT (white), SB-mNG (green), SB^P55A^-mNG (red), SB^P55,92A^-mNG (yellow) and SB^P55,92,94A^-mNG (magenta). Error bars indicate standard deviations, with dotted lines denote the mean. Different letters indicate statistically significant differences between groups (ANOVA with post-hoc Tukey HSD, *α*=0.05). Colony size of SB glycan variants reduces progressively as number of mutated sites increased. (G) Scatter dot plot showing the number of peripheral gametophores after 21 days of growth on AT medium. Genotypes, error bars, and statistical analysis are as described in (F). Numbers of gametophore are different in SB glycan variants.

Next, we phenotypically characterized these SB glycosylation variants with reference to WT and SB-mNG. Consistent with our previous observation that extremely high SB expression impaired growth and gametophore formation (Teh et al., 2022), moderate SB-mNG overexpression in the *sb sbl* background reduced the colony size (Fig. S1C-E). These lines were therefore unsuitable for subcellular localization analyses, as excessive ectopic expression frequently leads to protein mislocalization (Grefen et al., 2010). This growth inhibition was not attributable to the mNG tag itself, because expression of untagged SB cDNA produced similar defects (Teh et al., 2022). Therefore, further experiments were conducted with a carefully selected SB-mNG line that was almost phenotypically indistinguishable from WT, in terms of colony size and growth morphology (Fig. 1E, F).

When grown on AT medium, which restricts chloronema-to-caulonema transition, colony growth progressively decreased from SB^P55A^-mNG to SB^P55,92A^-mNG and SB^P55,92,94A^-mNG (Fig. 1E, F), indicating that hypoglycosylation compromises tip growth. These observations suggest that glycosylations at least at one or more of these sites, particularly P55, are required to sustain colony growth. In contrast, gametophore formation did not follow the same trend. On BCD medium, which promotes the chloronema-to-caulonema transition and thus gametohore development, the fewest gametophores were produced in SB-mNG and SB^P55A^-mNG (Fig. 1E, G). The reduction was less pronounced in SB^P55,92A^-mNG and SB^P55,92, 94A^-mNG (Fig. 1E, G), suggesting that glycosylations at these sites exerts an inhibitory effect on gametophore formation. This is consistent with the fully glycosylated SB-mNG showing the strongest inhibition and indicates that glycosylation at P55 is not essential for this inhibitory activity. On the other hand, glycosylation at P92 and P94 appears to contribute partially, in agreement with our earlier conclusion that additional glycosylation sites exist in SB. Together, our data indicate that SB plays an inhibitory role in colony growth and gametophore formation, with differential contributions from glycosylations at P55, P92 and P94. Because SB^P55,92, 94A^-mNG includes all three mutated glycosylation sites and exhibited the most severe growth inhibition in colony size, we selected this variant for further characterization.

### SB glycosylation is required for its intracellular trafficking

Glycosylation of AGPs has been implicated in their intracelluar trafficking, particularly in efficient secretion (Xue et al., 2017; Zhang et al., 2019). To determine whether the hypoglycosylated SB^P55,92, 94A^-mNG variant exhibits any trafficking defects, we compared its subcellular localization with that of SB-mNG. SB-mNG localised to punctate structures, some of which partially colocalised with the endocytic tracer FM4-64 (Fig. 2A), indicating that a sub population of SB undergoes endocytosis via the *trans*-Golgi network (TGN). In contrast, SB^P55,92, 94A^-mNG displayed a more diffusive distribution with significantly fewer puncta (Fig. 2B, C). Because SB is a membrane protein and should not appear cytoplasmic, the diffuse signal raised the possibility of increased free mNG generated by cryptic cleavage of SB fusion proteins. To test this, we quantified the abundance of full-length fusion proteins and free mNG using protein fractionation assay followed by mNG immunoblotting. In membrane fractions (P_100_), SB^P55,92, 94A^-mNG fusion was more abundant than SB-mNG (Fig. 2D), ruling out reduced expression as the cause of fewer puncta. In soluble fractions (S_100_), free mNG levels were comparable between the two lines, indicating that the diffuse signal in SB^P55,92, 94A^-mNG cannot be attributed to increased release of free mNG.

**Figure 2.**
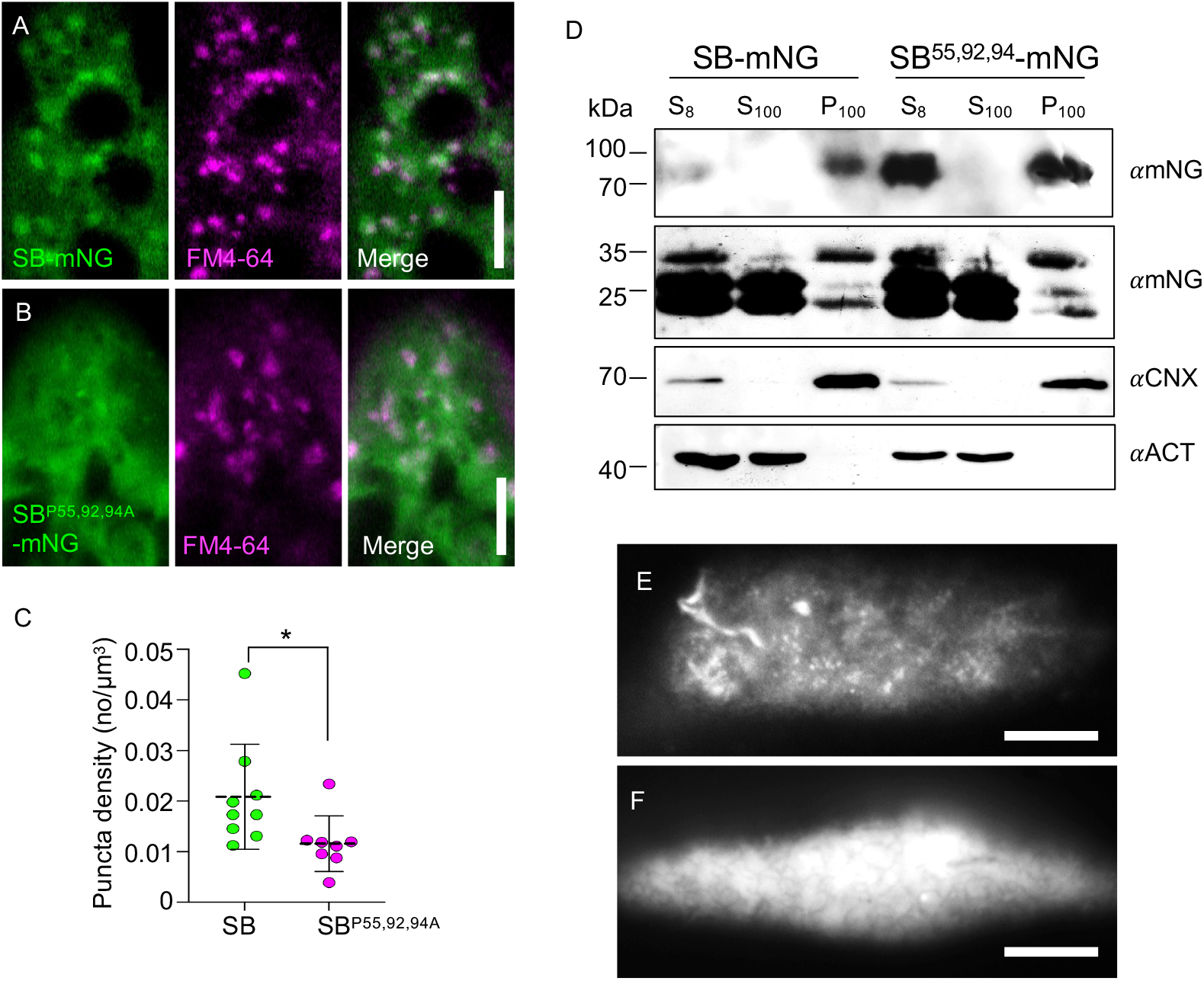
A subpopulation of SB-mNG undergoes endocytosis. (A, B) Confocal images of apical protonemal cells expressing SB-mNG (A) or SB^P55,92,94A^-mNG (B). Images are maximum-intensity of z-stacks. mNG and FM4-64 signals are shown in green and magenta, respectively. Scale bar, 5 µm. A subset of SB-mNG puncta, but not SB^P55,92,94A^-mNG puncta, colocalized with FM4-64. (C) Scatter plot showing puncta density of SB-mNG (green circles) and SB^P55,92,94A^-mNG (magenta circles). Staistical signifiance was assessed using a two-tailed Mann-Whitney test, * (P≤0.05). Puncta density of SB^P55,92,94A^-mNG was reduced. (D) Immunoblot of soluble (S_100_) and membrane (P_100_) fractions isolated from SB-mNG and SB^P55,92,94A^-mNG lines following ultracentrifugation at 100, 000 *g*. SB fusion proteins were detected using an anti-mNG antibody (*α*mNG). Calnexin (*α*CNX) and Actin (*α*ACT) were used as membrane and cytosolic markers, respectively, to assess cross contaminations between fractions. SB^P55,92,94A^-mNG accumulates at higher levels than SB-mNG, while free mNG levels are comparable between lines. (E, F) TIRF microscopic images of apical protonemata from expressing SB-mNG (E) or SB^P55,92,94A^-mNG (F). Scale bar, 5 µm. SB-labelled foci were observed in SB-mNG but not SB^P55,92,94A^-mNG.

Because only a subset of SB-mNG puncta colocalised with FM4-64, we hypothesized that the remaining SB-mNG puncta correspond to plasma membrane-associated signal. To clarify this point, we used total internal reflection fluorescence (TIRF) microscopy to examine the spatial distribution of SB-mNG and SB^P55,92, 94A^-mNG near the plasma membrane. In SB-mNG lines, TIRF imaging revealed dynamic and discrete SB-mNG-labelled foci moving along the plasma membrane against a weak fluorescent background (Fig. 2E, Fig. S2A). In contrast, such foci were not detected in SB^P55,92, 94A^-mNG, where the plasma membrane was instead marked by a strong and relatively uniform fluorescent signal (Fig. 2F, Fig. S2B). Notably, mNG signal was present at the plasma membrane in both genotypes, indicating that SB association with the plasma membrane is not dependent on glycosylation. However, the absence of discrete foci in SB^P55,92, 94A^-mNG suggests that glycosylation at P55, P92, and P94 may be required for the formation or stabilization of SB-mNG labelled foci at the plasma membrane.

### Vacuolar trafficking is affected in hypoglycosylated SB

Since a subset of SB colocalised with FM4-64 (Fig. 2A), we next investigated the trafficking route(s) taken by SB after endocytosis. We first tested whether SB undergoes endosomal recycling by treating SB marker lines with Brefeldin A (BFA), a fungal metabolite that inhibits the GNOM ARF-GEF-mediated endosomal recycling pathway(Geldner et al., 2003). Neither SB-mNG nor SB^P55,92, 94A^-mNG accumulated in the FM4-64-labelled BFA compartments formed by TGN agglomerations (Fig. S3A, B), indicating that endocytosed SB does not recycle back to plasma membrane through TGN. Notably, SB-mNG was detected on spherical, FM4-64-stained membranous structures which we interpret as intravacuolar membranes or the tonoplast (Fig. 3A). Quantification of the FM4-64 and SB-mNG fluorescence intensities along these structures showed near-perfect alignment of intensity peaks (Fig. 3B), confirming their colocalisation. These results support the conclusion that endocytosed SB is targeted to the vacuole. In contrast, hypoglycosylated SB^P55,92, 94A^-mNG showed no detectable association with these internal membranes (Fig. 3C), and fluorescence intensity profiling revealed no overlapping peaks with FM4-64 (Fig. 3D). To further confirm that endocytosed SB is trafficked to the vacuole, we disrupted vacuolar transport pathways using Wortmannin, an inhibitor of phosphatidylinositol 3-kinases (PI3K) signalling that induces homotypic fusion of pre-vacuolar compartments (PVC) in Arabidopsis (Wang et al., 2009; Munch et al., 2015). Upon Wortmannin treatment, SB-mNG accumulated on MDY-64-labelled (a vacuolar membrane dye(Radin et al., 2021)) membranous structures (Fig. 3E, F), confirming that SB is sorted to the vacuole following endocytosis. In contrast, Wortmannin-treated SB^P55,92, 94A^-mNG exhibited predominantly diffuse fluorescence and remained adjacent to, but not colocalized with, the MDY-64-stained tonoplast, as reflected by the lack of overlapping intensity peaks (Fig. 3G, H). Together, these results demonstrate that vacuolar trafficking of SB requires full glycosylation at P55, P92, and P94.

**Figure 3.**
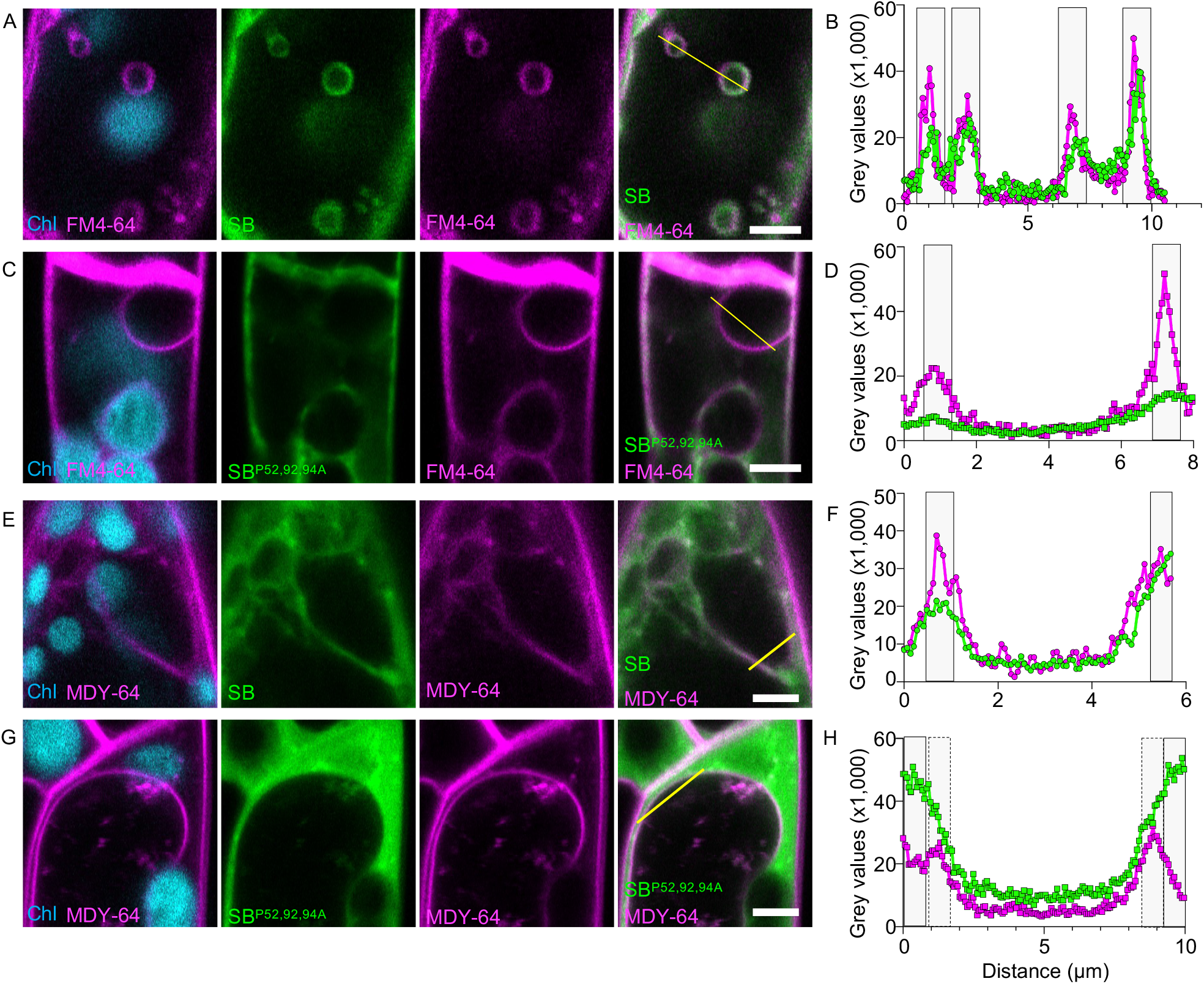
Endocytosed SB-mNG is trafficked to the vacuole. (A-D) Confocal images of apical protonemal cells expressing SB-mNG (A, B) or SB^P55,92,94A^-mNG (C, D) following treatment with 50 µM BFA. Images are single optical sections. Chloroplast autofluorescence, mNG, and FM4-64 signals are shown in blue, green, and magenta respectively. Yellow lines indicate regions used for fluorescence intensity profiling in (B) and (D). Scale bar, 5 µm. SB-mNG, but not SB^P55,92,94A^-mNG, localizes to the vacuolar membrane following BFA treament. (E-H) Confocal images of apical protonemal cells expressing SB-mNG (E, F) or SB^P55,92,94A^-mNG (G, H) following treatment with 33 µM Wmn. Images are single optical sections. Chloroplast autofluorescence, mNG, and FM4-64 signals are shown in blue, green, and magenta respectively. Yellow lines indicate regions used for fluorescence intensity profiling in (F) and (H). Scale bar, 5 µm. SB-mNG, but not SB^P55,92,94A^-mNG, localizes to vacuolar membrane following Wmn treament.

### SB is required to maintain cell wall integrity but not cell wall mechanics

We previously showed that galactose content in the *sb sbl* cell wall is markedly reduced compared with WT, suggesting a role for SB in the cell wall remodelling or biosynthesis (Teh et al., 2022). To directly link SB function, and its glycosylation status to cell wall architecture, we examined cell wall ultrastructure in *sb sbl*, SB-mNG, and SB^P55,92, 94A^-mNG lines alongside WT controls using transmission electron microscopy (TEM). In WT, the polysaccharide-rich primary cell wall appears as a moderately electron-opaque layer above the plasma membrane (Fig. 4A) and is topped by a ∼30 nm-thick cuticular layer that separates the wall from the cuticle proper (Yeats and Rose, 2013). In contrast, most *sb sbl* TEM micrographs (9/13) lacked this distinct cuticular layer; instead we observed a disintegrated and irregular layer containing sparse cutin-like materials (Fig. 4A, Fig. S4A). The *sb sbl* cell walls are also thicker (Fig. 4A) and displayed prominent mis-oriented speckles, further indicating structural disruption (Fig. 4A). Both SB-mNG and SB^P55,92, 94A^-mNG retained an identifiable cuticular layer (Fig. 4A), although quantitative measurements did not indicate a significant difference in their thickness (Fig. S4B).

**Figure 4.**
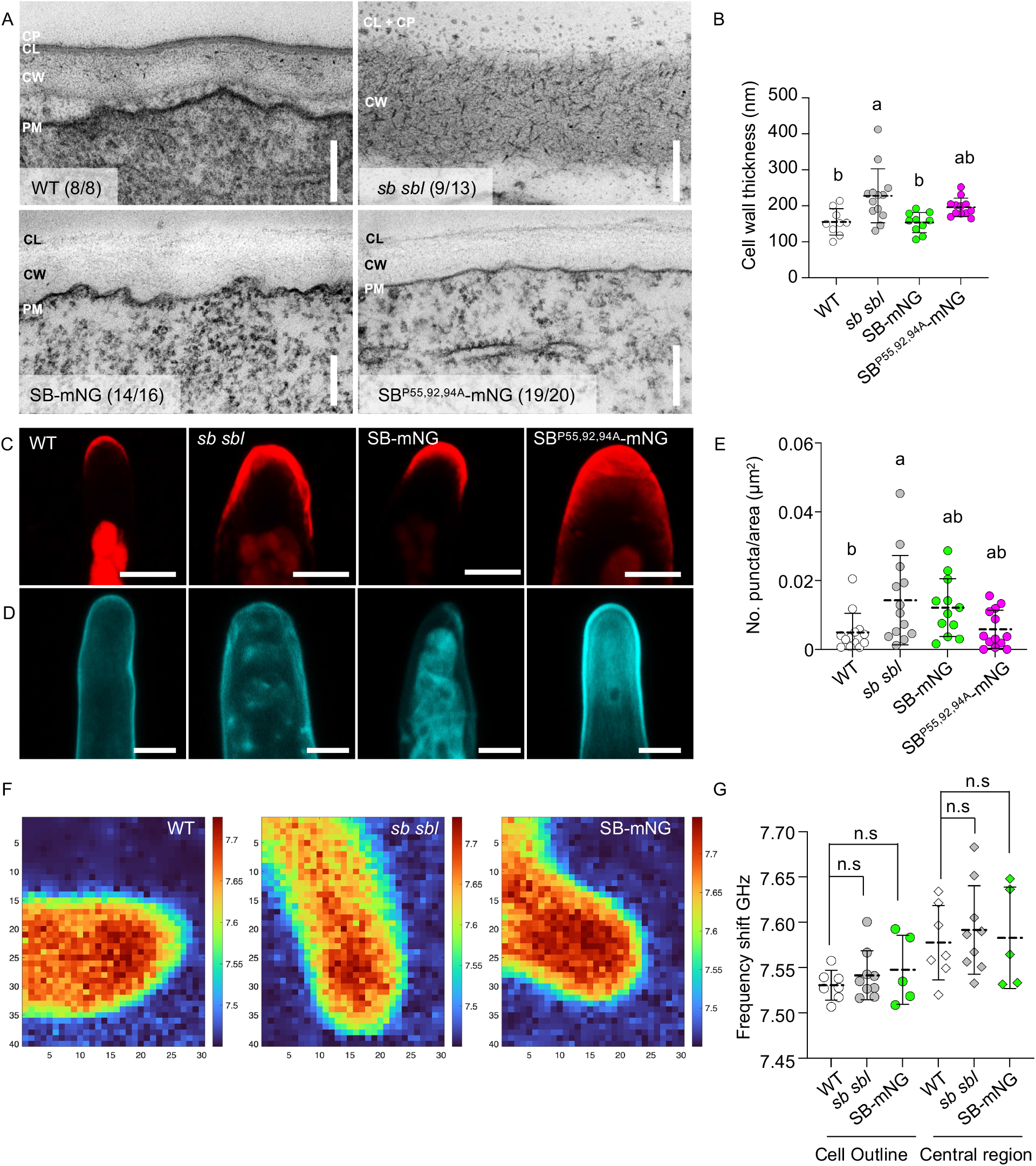
Cell wall integrity is disrupted in SB alleles. (A) Representative TEM micrographs of WT (top left), *sb sbl* (top right), SB-mNG (bottom left), and SB^P55,92,94A^-mNG (bottom right). Numbers in parentheses indicate the fraction of micrographs displaying the corresponding ultrastructural morphology.. CP, cuticle proper; CL, cuticular later; CW, cell wall; PM, plasma membrane. Scale bar, 200 nm. Cell wall ultrastructure in *sb sbl* is altered. (B) Scatter dot plot showing cell wall thickness in WT (white circles), *sb sbl* (grey circles), SB-mNG (green circles), and SB^P55,92,94A^-mNG (mangenta circles). Error bars indicate standard deviation. Different letters denote statistically significant differences between groups (one-way ANOVA with Tukey’s HSD test, *α*=0.05). Cell wall in *sb sbl* is thicker. (C, D) Confocal images of protonemata labelled with LM2 antibody (C) or stained with Calcoflour White (D). Images are sum projections of z-stacks. Scale bar, 10 µm. Calcofluor-white staining in *sb sbl* is altered. (E) Scatter plot showing the density of Calcofluor White-stained punctate in WT (white), *sb sbl* (grey), SB-mNG (green) and SB^P55,92,94A^-mNG (magenta). Statistical significance was assessed using a two-tailed Mann-Whitney test (*P≤0.05). No. of puncta per area size is increased in *sb sbl*. (F) Representative heatmaps of frequency shifts measured in apical tip cells. (G) Quantification of Brillouin frequency shifts in WT (white), sb sbl (grey), and SB-mNG (green), measured at the cell periphery (circles) and central regions (diamonds). Statistical analyss was performed using one-way ANOVA with Šidák’s multiple-comparison test (*α*=0.05). n.s., not significant. Cell wall stiffness was not changed in SB alleles.

To further assess the relationship between SB and cell wall compositions, we next compared cell wall polysaccharide spatial distribution among SB alleles. Immunostaining with LM2, which recognizes AGPs (Smallwood et al., 1996) revealed the previously reported tip-enriched pattern in WT (Lee et al., 2005), which was unchanged in all SB alleles (Fig. 4C), indicating that SB does not affect the localization or abundance of other AGPs. Similarly, staining for unesterified pectins using LM19 (Verhertbruggen et al, 2009) showed uniform distribution across all genotypes (Fig. S4C), suggesting that SB function does not directly regulate pectin composition. We next examined the cellulose microfibril organization using calcofluor-white, a fluorescent dye that binds to β-1,4 glycosidic bonds of cellulose. In WT, calcofluor-white staining was smooth and evenly distributed along the cell wall (Fig. 4D). In contrast, *sb sbl* exhibited a marked increase in irregular puncta per unit area (Fig. 4C, E), indicative of disrupted cellulose microfibril distribution or organization. Notably, this staining pattern was also observed in the SB-mNG and hypoglycosylated SB^P55,92, 94A^-mNG, although with a moderate increment (Fig. 4E).

In plants, cell wall integrity is a key mechanism for monitoring and maintaining the structural and mechanical properties of the cell wall(Vaahtera et al., 2019; Bacete et al., 2022). Perturbations in cell wall integrity are often associated with changes in cell wall viscoelasticity, which can be quantified using Brillouin light scattering(Samalova et al., 2023; Alonso Baez et al., 2026). Because ultrastructural and immunostaining analyses suggested that cell wall integrity is compromised in *sb sbl*, we therefore investigated whether cell wall mechanical properties are altered in SB-related genotypes (*sb sbl* and SB-mNG) using Brillouin light scattering. Brillouin frequency shifts and linewidths were measured at both the outline (cell walls) and central regions (cytoplasm and cell walls) of tip cells (Fig. S4D). No pronounced differences were detected in Brillouin frequency shifts, which reflect stiffness (Fig. 4F, G), or in linewidths, which are indicative of viscosity (Fig. S4E, F), across the genotypes examined. Based on these observation, we did not further assess cell wall mechanical properties in SB^P55,92, 94A^-mNG lines, and conclude that disruption of SB does not measurably affect cell wall mechanical properties.

### SB alleles are hypersensitive to cell wall stress

To further test the functional link between SB and cell wall integrity, we examined the sensitivity of SB alleles to cellulose biosynthesis inhibitor isoxaben (ISX) and 2,6-dichlorobenzonitrile (DCB). Unlike ISX which specifically inhibits cellulose synthase (CESA), DCB inhibits both CESA and cellulose synthase-like D (CSLD) enzymes, thereby disrupting fibrillar β-1,4-glucan synthesis and compromising cell wall integrity. We reasoned that SB alleles with altered cellulose organization would display increased sensitivity to cell wall stress induced by cellulose biosynthesis inhibitors. Under mock conditions, SB^P55,92, 94A^-mNG exhibited reduced growth relative to WT (Fig. 5A left panel, Fig. 5B), whereas *sb sbl* formed significantly larger colonies (Fig. 5A, B), consistent with previous observation (Fig. S1C). Upon 5 µM DCB treament, WT plants showed a marked reduction in colony size after 10 days, an indication of impaired tip growth, as reported previously (Favery et al., 2001; Wu et al., 2023). This growth inhibition was significantly exacerbated in *sb sbl*, SB-mNG and SB^P55,92, 94A^-mNG, all of which exhibited smaller colonies than DCB-treated WT (Fig. 5A right panel, Fig. 5B). These results demonstrate that disruption of SB function enhances sensitivity to DCB-induced cell wall stress. Notably, DCB hypersensitivity was partially alleviated in the hypoglycosylated SB^P55,92, 94A^-mNG line compared with SB-mNG (Fig. 5A right panel, Fig. 5B), further highlighting the contribution of SB glycosylation to its role in cell wall integrity. In contrast, treatment with 20 µM ISX had only a mild effect on the colony size across all genotypes (Fig. S5A, B), consistent with previous reports that CSLD, rather than CESA, is the primary cellulose synthase enzymes in *P. patens* (Roberts et al., 2024). Together, these findings establish SB and its proper glycosylation as a critical determinant of cell wall integrity, possibly through regulation of cellulose microfibril organization.

**Figure 5.**
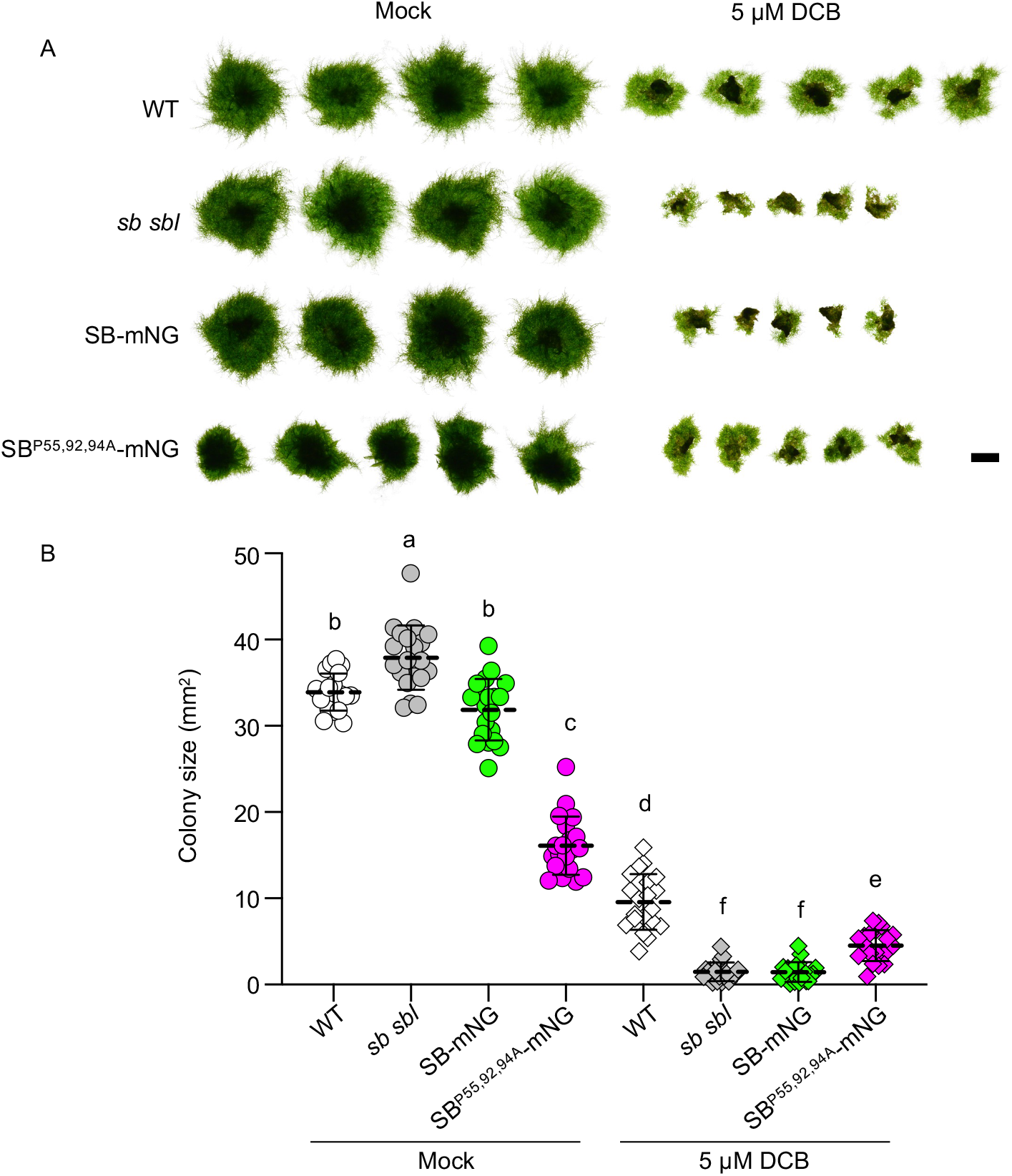
SB alleles are hypersensitive to cell wall stress. (A) Representative images of10-day-old colonies of WT, sb sbl, SB-mNG, and SB^P55,92,94A^-mNG under mock conditions (left) or in the presence of 5 µM DCB (right). Scale bar, 2mm. Colonies from the same experiment were imaged individually and stitched together. All SB alleles exhibited hypersensitivity to DCB treatment. (B) Scatter plot showing colony size of mock-treated (circles) and 5 µM DCB-treated (diamond) plants. Different indicate statistically significant differences between groups (ANOVA with Tukey’S HSD test, *α*=0.05). Although all SB alleles are hypersensitive to DCB, the hypersensitivity is alleviated in SB^P55,92,94A^-mNG.

### Hypoglycosylated SB fails to localise on cross wall during tip growth

To directly assess whether protonemal tip growth was altered/compromised in SB mutan and overexpression lines with altered colony size, we performed time-lapse imaging of 7-day-old protonemata. Consistent with the increased colony size of *sb sbl*, protonemal tip growth speed was significantly higher in *sb sbl* compared with WT (WT: 0.197 ± 0.046 µm/min; *sb sbl*: 0.265 ± 0.068 µm/min, Fig. 6B), resulting in longer protonemata at synchronized time points (Fig. 6A). It is possible that compromised cell wall integrity in *sb sbl* enabled faster cellular expansion. In contrast, tip growth speed were reduced in SB-mNG and SB^P55,92, 94A^-mNG (0.168 ± 0.065 µm/min and 0.134 ± 0.021 µm/min respectively; Fig. 6B) consistent with their smaller colony sizes (Figs 1F, 5A).

**Figure 6.**
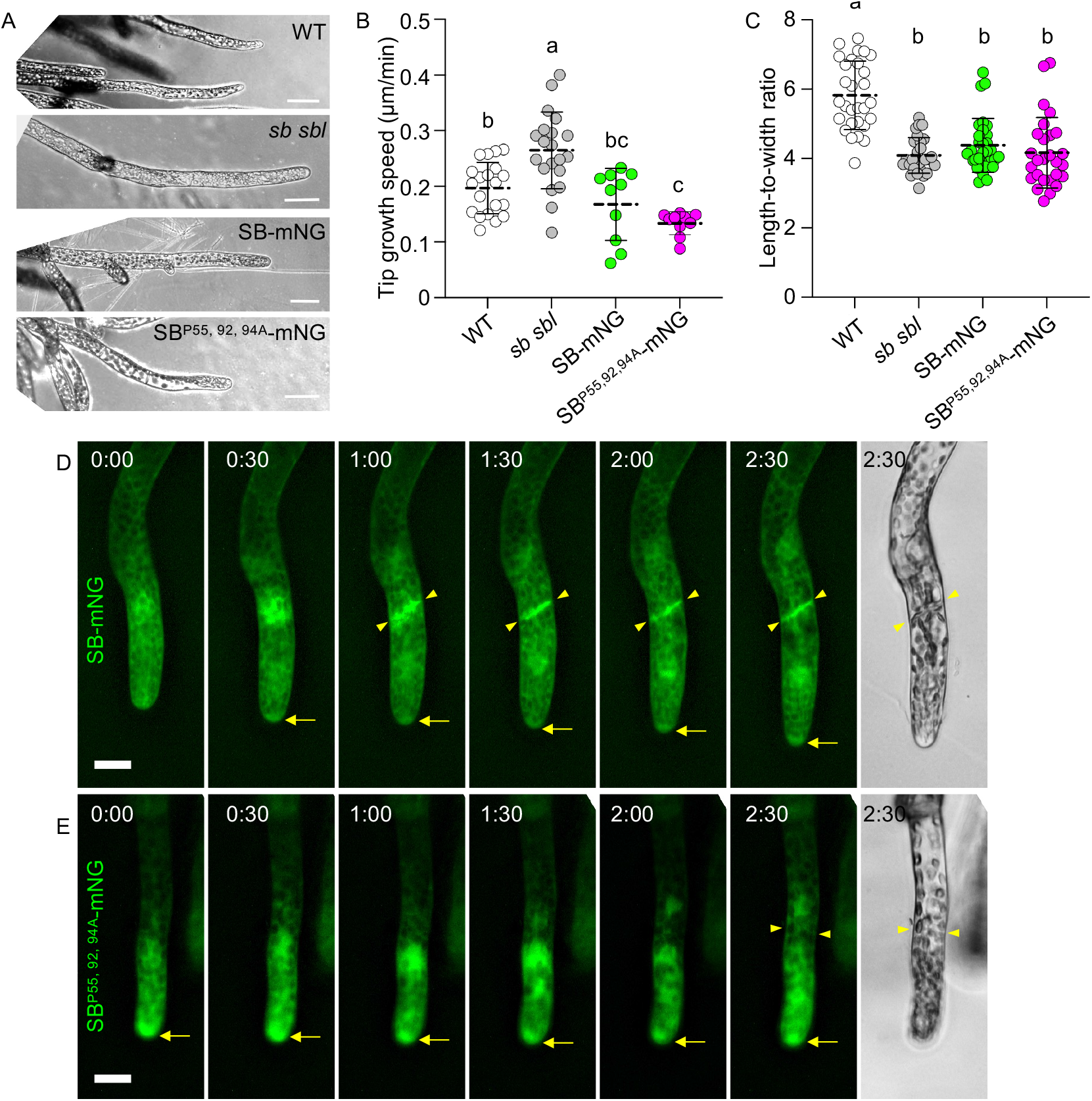
SB glycosylation is required for apical tip growth and localization to nascent cross walls. (A) Representative snapshots images of protonemal filaments after 17 h 30 min growth. Scale bar, 50 µm. Protonemata of *sb sbl* are longer compared to other genotypes. (B, C) Scatter plot showing apical tip growth rates (B) and length-to-width ratio (C) of WT (white circles), *sb sbl* (grey circles), SB-mNG (green circles) and SB^P55,92,94A^-mNG (magenta circles). Different letters indicate statistically significant differences between groups (one-way ANOVA with Tukey’s multiple-comparison test, *α*=0.05). Elongation speed is the fastest and slowest in *sb sbl* and SB^P55,92,94A^-mNG, respectively. (D, E) Representative snapshots images from time-lapse imaging of SB-mNG (D) and SB^P55,92,94A^-mNG (E). Time stamps (top left) are synchronized to the end of mitosis. Arrow heads indicate newly formed cross walls, and arrows indicate the apical tip. Scale bar, 20 µm. SB-mNG, but not SB^P55,92,94A^-mNG, localised on crosswalls.

Because tip growth is a highly polarized cell elongation process and contributes to the elongated and cylindrical morphology of protonemata, we next examined whether altered tip growth speed affected cellular dimensions/geometry. Protonemal cell length was similar among WT (105.66 ± 11.97 µm), *sb sbl* (101.43 ± 12.35 µm), and SB-mNG (103.60 ± 19.15 µm), whereas SB^P55,92, 94A^-mNG exhibited greater variability and a reduced mean cell length (82.34 ± 17.16 µm) (Fig. S6A). In contrast, cell width differed markedly among genotypes: *sb sbl* (24.98 ± 3.15 µm) and SB-mNG (24.02 ± 4.63 µm) were significantly wider than WT (18.46 ± 2.79 µm) and SB^P55,92, 94A^-mNG (20.02 ± 2.30 µm, Fig. S6B). Collectively, these changes in cell length and width resulted in a reduced length-to-width ratio across all SB alleles (WT: 5.82 ± 0.99; *sb sbl*: 4.09 ± 0.51; SB-mNG: 4.31 ± 0.78; SB^P55,92, 94A^-mNG: 4.17 ± 1.02; Fig. 6C), demonstrating that disruption of SB function alters the cellular geometry, consistent with impaired cell wall integrity and perturbed tip growth regulation.

Since SB intracellular trafficking depends on its glycosylation (Fig. 2, 3), we next examined whether the subcellular localisation of hypoglycosylated SB^P55,92, 94A^-mNG is affected during protonemal tip growth. Time-lapse imaging of SB-mNG revealed a distinct and persistent polar enrichment at the growing tip (arrow, Figure 6D, t=0:30 to t=2:30), corresponding to the apical “clear zone”, which is enriched with Golgi, ER and vesicles(Bove et al., 2008; Furt et al., 2012). Notably, SB-mNG also localised to nascent cross walls as early as ∼2 h before completion of mitosis (arrow head, Figure 6D, t=1:00). SB-mNG localisation on the cross walls is also observed in gametophore divisions (Fig. S6C). By contrast, although SB^P55,92, 94A^-mNG also accumulated at the apical “clear zone” during tip growth (arrow, Figure 6E, t=0:00 to t=2.30), it failed to localize to newly formed cross walls, even when cross wall formation was clearly evident at the end of mitosis (arrow head, Figure 6E, t=2:30). It is important to note that SB^P55,92, 94A^-mNG localization on plasma membrane was eventually established at a much later time point (Fig. S6C), consistent with observation using TIRF imaging (Fig. 2E,F). This selective and transient loss of cross wall localization indicates that efficient secretion and/or retention of SB at the plasma membrane during cell division requires full glycosylations at P55, 92 and 94.

### Functional conservation of SB in tip growth

Carbohydrate-active enzyme (CAZyme) families, including those responsible for AGP glycosylation, play central roles in the synthesis and modifications of cell wall polysaccharides. These families originated in streptophyte algae Zygnematophyceae, the closest relatives of land plants, and lineage-specific CAZyme expansions have been proposed to underpin cell wall innovations required for terrestrialization (Feng et al., 2024). Given that SB glycosylation regulates its intracellular trafficking and cross-wall localization to control tip growth, we next asked whether SB function is conserved in vascular plants. We first ectopically expressed a 35S promoter-driven, YFP-tagged SB in Arabidopsis (*35S:SB-YFP*). Although rosette growth was largely unaffected, inflorescence length was moderately reduced compared with WT and a control line expressing free YFP (*35S:YFP*) (Fig. S7A, B). To assess whether SB was properly processed in this heterologous context, we examined its glycosylation status. Immunoblot analysis of membrane protein fractions (Lane P in Fig. S7C) revealed a high-molecular weight smear extending beyond 100 kDa, distinct from the SB profile observed in *P. patens* (Fig. 1C), suggesting that SB may have been incorrectly glycosylated when expressed in a heterologous system.

To circumvent this, we confined SB expression to a defined tip-growing cell type. SB-mNG was expressed specifically in root hairs using the trichoblast-specific AtEXP7 promoter (Cho and Cosgrove, 2002). Stable Arabidopsis lines expressing AtEXP7_pro_:SB-mNG (#4) displayed pronounced root hair-specific fluorescence and produced short, stunted root hairs (Fig. 7A, B), indicative of inhibited tip growth, similar to the phenotype observed in *P. patens*. To exclude developmental staging effects, root hair length was quantified in the third (#3) to fifth (#5) differentiated root hairs. We observed significantly shorter root hair length (#3: 0.034 ± 0.007 mm, #4: 0.038 ± 0.010 mm, #5: 0.034 ± 0.007 mm; Fig. 7C) than those of WT (#3: 0.119 ± 0.053 mm, #4: 0.158 ± 0.051 mm, #5: 0.187 ± 0.060 mm; Fig. 7C), demonstrating that SB overexpression inhibits root hair elongation early after protrusion. We next tested whether this inhibitory effect depends on SB glycosylation. Stable lines expressing the hypoglycosylated variant AtEXP7_pro_: SB^P55,92, 94A^-mNG were generated in the same manner. In contrast to SB-mNG, expression of SB^P55,92, 94A^-mNG resulted in a partial inhibition of root hair length (#3: 0.097 ± 0.118 mm, #4: 0.077 ± 0.056 mm, #5: 0.087 ± 0.061 mm; Fig. 7C), indicating alleviation of the tip growth defect. Together, these results demonstrate that SB-mediated regulation of tip growth is glycosylation-dependent and suggest that this mechanism is at least partially conserved between bryophytes and vascular plants.

**Figure 7.**
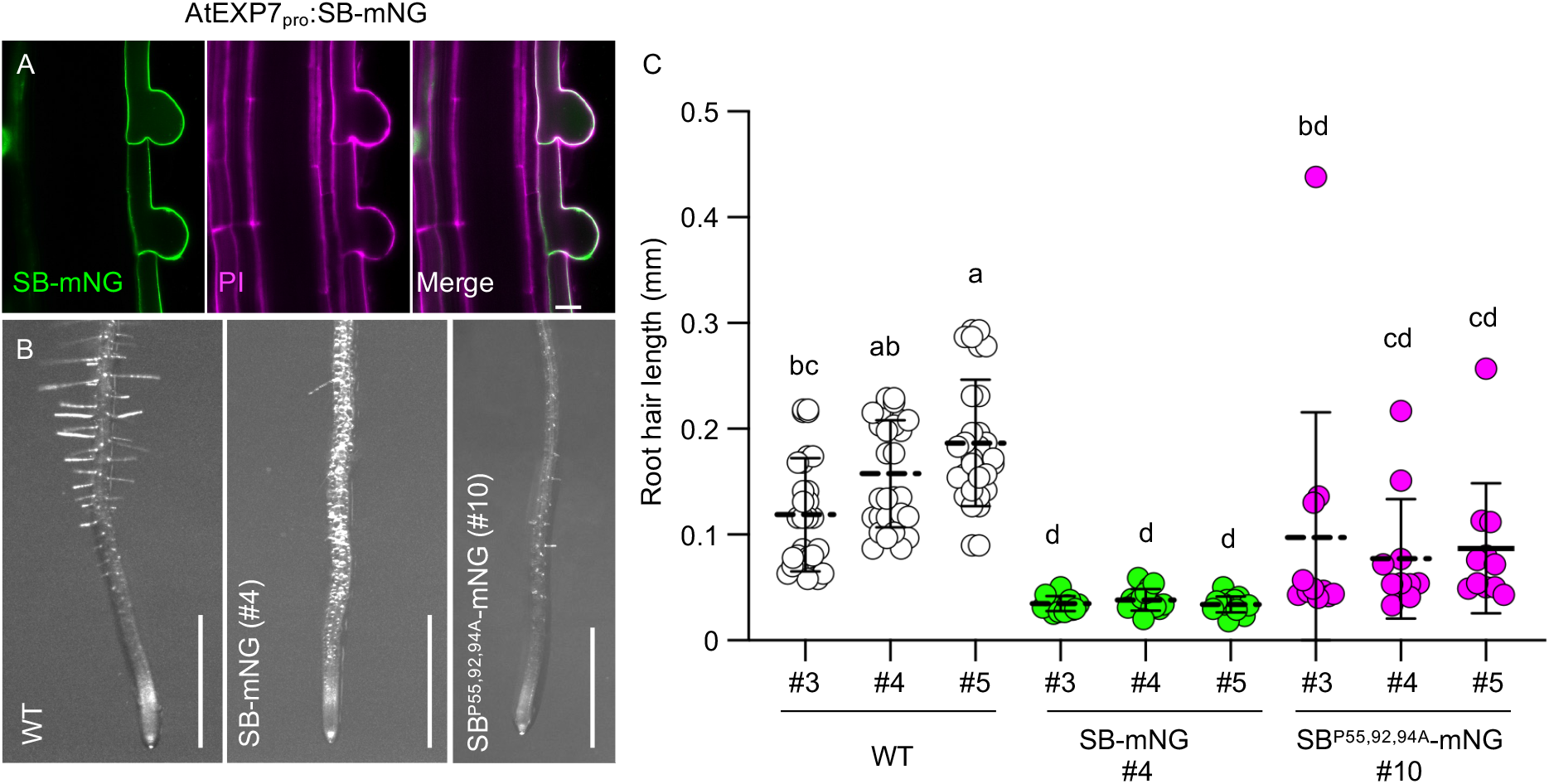
SB function in tip growth is conserved in Arabidopsis. (A) Confocal images of root hairs from 5-day-old Arabidopsis seedlings expressing AtEXP7_pro_:SB-mNG (green) and stained with propidium iodide (PI, magenta). Images are maximum-intensity projections of z-stack. Scale bar, 20 µm. AtEXP7_pro_-driven SB-mNG expression is restricted to root hair. (B) Bright-field images of root tissues from 5-day-old WT, AtEXP7_pro_:SB-mNG and AtEXP7_pro_: SB^P55,92,94A^-mNG seedlings. Scale bar, 1 mm. Root hair tip growth is inhibited in AtEXP7_pro_:SB-mNG and AtEXP7_pro_: SB^P55,92,94A^-mNG. (C) Scatter plot showing root hair length in WT (white circles), AtEXP7_pro_:SB-mNG (green circles) and AtEXP7_pro_: SB^P55,92,94A^-mNG (magenta circles). Different letters indicate statistically significant differences between groups (one-way ANOVA with Tukey’s multiple-comparison test, *α*=0.05). Root hair inhibition is partially rescued in AtEXP7_pro_: SB^P55,92,94A^-mNG.

## Discussion

Since their initial isolation from suspension-cultured sycamore cells in 1969, AGPs have been recognized as ubiquitous components of all land plants and implicated in a wide range of developmental and physiological processes (see excellent reviews by (Tan et al., 2012; Silva et al., 2020; Hromadova et al., 2021; Ma and Johnson, 2023)). Despite their abundance, the molecular mechanism by which AGPs function within the cell wall, as well as their downstream interactors, remains poorly understood. In *P. patens*, early evidence for a structural role of AGPs in the cell wall came from the inhibition of apical tip growth following treatment with the AGP-binding reagent β-Yariv (Lee et al., 2005), as well as reduced protonemal growth in an *agp1* knockout mutant (Lee et al., 2005). Here, through detailed characterisation of SB loss-of-function mutant (*sb sbl*) and semi-overexpressors (SB-mNG, SB^P55,92, 94A^-mNG), we establish SB as a functionally important AGP required for cell wall integrity and regulating apical tip growth. Importantly, we demonstrate that three hydroxyproline glycosylation sites are critical for SB intracellular trafficking and activity.

### Biological relevance of SB intracellular trafficking

Secretion of cell wall matrix components – including polysaccharides, glycoproteins, and proteoglycans - is crucial for apical tip growth. In tip-growing cells, secretory organelles such as endoplasmic reticulum, Golgi apparatus, and secretory vesicles are asymmetrically enriched within the apical “clear zone” (Kim and Brandizzi, 2016; Bibeau et al., 2021). Although AGP secretion to the plasma membrane remains poorly characterized, GPI-anchored AGPs are thought to follow non-conventioal, concentrative sorting pathway (Borner et al., 2005). In Arabidopsis pollen tube, the actin-nucleating protein FORMIN5 recruits and deposits AGP23 into the cell wall, highlighting the importance of precisely timed AGP secretion for cell wall compositions and integrity during polarized growth (Li et al., 2024).

Our data show that hypoglycosylated SB^P55,92, 94A^-mNG is mis-sorted, exhibiting aberrant vacuolar targeting (Fig. 3) and impaired secretion to the plasma membrane during apical tip growth (Fig. 6E). These observations suggest that proper glycosylation is required for accurate SB sorting at the TGN, where secretory and endocytic pathways converge. In plants, type II AG biosynthesis occurs predominantly in the medial- to *trans*-Golgi (Parsons et al., 2019), and Hyp-linked arabinogalactan modifications have been shown to enhance protein secretion efficiency (Xue et al., 2017; Zhang et al., 2019). One plausible model is that Hyp-arabinoxylation promotes correct SB folding and serves as a quality-control checkpoint prior to packaging into secretory vesicles. Alternatively, or additionally, these glycan moieties may function as a sorting signal at the TGN, preventing SB from being misdirected into vacuolar degradation pathways.

Although all AGPs possess an N-terminal signal peptide and are therefore secreted by default, it remains unclear whether secreted AGPs generally undergo endocytic uptake. The Arabidopsis Fasciclin-like AGP FLA4 undergoes endosomal recycling following internalization (Xue et al., 2017), whereas endocytosed SB is trafficked to the vacuole rather than recycled (Fig. 3). The biological significance of SB vacuolar targeting is not yet clear; however, we speculate that this may represent a regulatory mechanism to maintain low SB abundance at the cell surface, thereby fine-tuning cell wall extensibility during tip growth. Consistent with this idea, endogenous SB expression is extremely low, and SB overexpression leads to severe growth arrest (Teh et al., 2022). Alternatively, SB may function as a cargo receptor in clathrin-mediated endocytosis, as has been shown in the uptake of rare earth elements (Wang et al., 2019).

### Mechanistic role of SB in cell wall integrity

Ultrastructural analysis (Fig. 4A), immunolocalisation (Fig. 4B) and pharmacological perturbations (Fig. 5) collectively reveal a central role for SB in maintaining cell wall integrity. In the *sb sbl* amorphic mutant, the most pronounced defects include loss of the cuticular layer and increased cell wall thickness (Fig. 4A), likely reflecting compensatory responses to compromised wall integrity. These defects correlate with disorganized cellulose microfibril patterns (Fig. 4C), implicating SB in cellulose-related wall architecture. Notably, cell wall mechanical properties remained largely unchanged upon SB disruption (Fig. 4F, G), suggesting that, in this context, cell wall integrity can be uncoupled from bulk cell wall mechanics. This observation parallels previous findings that mechanosensing by the cell wall integrity sensor FERONIA is separable from the downstream cell wall reinforcement mediated by microtubule-guided cellulose biosynthesis (Malivert et al., 2021).

How does a surface-localised SB influence cell wall integrity? Two models have been proposed for AGP function. First, AGP may directly influence cellulose synthesis or assembly through physical interactions or cross-linking to other cell wall components (Hijazi et al., 2014; Lin et al., 2022). Second, AGPs may act as apoplastic Ca^2+^ capacitors, binding and releasing Ca^2+^ in a pH-dependent manner to regulate wall mechanics and signalling (Lamport et al., 2014). Currently, strong genetics and biochemical evidence supports the latter model: β-linked glucuronic acid residues in AGP reversibly bind Ca^2+^, and Arabidopsis mutants defective in β-glucuronyltransferase activity exhibit disrupted calcium waves as well as calcium-dependent developmental defects (Lamport and Varnai, 2013; Lopez-Hernandez et al., 2020).

Our findings link these models by showing that all SB alleles exhibit defects in cellulose organization (Fig. 4C) and hypersensitivity to the cellulose biosynthesis inhibitor DCB (Fig. 5), implicating SB in cellulose biosynthesis, assembly, or spatial organization. We propose that SB, as an intrinsically disordered and highly glycosylated protein (Fig. 1B), may function as a flexible scaffold that integrates/coordinates multimeric cellulose synthase complexes with matrix polysaccharides such as pectin to preserve wall integrity. Supporting this idea, covalent linkages between AGP, pectins, and hemicellulosic arabinoxylan have been described in a proteoglycan complex named ARABINOXYLAN PECTIN ARABINOGALACTAN PROTEIN (Tan et al., 2013). Moreover, tip growth defects in hydroxyproline *O*-arabinosyltransferase (HPAT) mutants can be rescued by exogenous cellulose application (MacAlister et al., 2016), further supporting a functional link between AGP glycosylation and cellulose deficiency. Given the extraordinary diversity of the AGP family, these mechanistic models are unlikely to be mutually exclusive. Instead, AGPs such as SB may integrate calcium signaling and cell wall structural organization - two interdependent processes essential for maintaining wall integrity during polarized tip growth.

The functional versatility of SB in cell wall integrity is reflected in the differential contributions of glycosylation at residues P55, P92, and P94. Whereas mutations at these three sites had an additive effect on apical tip growth (Fig. 1F), their impact on gametophore formation was qualitatively distinct (Fig. 1G), suggesting context-dependent roles for SB glycosylation during development. Because these glycosylation sites reside within the apoplastic, intrinsically disordered ectodomain of SB, we propose that they may infleunce the SB calcium-binding and, consequently, systemic calcium wave propagation during gametophore morphogenesis, particularly when SB is mislocalised (Fig. 2E,F, Fig. 3, Fig. 6E). In *P. patens*, calcium waves originate from the basal region of the gametophore and propagate directionally towards the apex (Storti et al., 2018). Inefficient secretion of hypoglycosylated SB to the plasma membrane during formative cell divisions could therefore perturb the signaling trajectory and compromise the transition to three-dimensional leaf shoot development. Future analysis comparing calcium wave dynamics in *sb sbl* and SB^P55,92, 94A^-mNG lines will be critical to directly test whether SB participates in calcium-mediated developmental signaling.

## Materials and methods

### Plant materials and growth conditions

*Physcomitrium patens* (Gransden isolate) was used as the wild-type (WT) throughout this study. Protonemal tissue was homogenized and cultured on BCDAT agar medium overlaid with cellophane (Shintech Instruments, Taiwan). Transgenic moss lines were generated by PEG-mediated transformation as described previously(Cove et al., 2009). Stable transformants were selected twice on BCDAT medium supplemented with appropriate antibiotics.

*Arabidopsis thaliana* ecotype Columbia (Col-0) was used as WT. Seeds were surface-sterilized and germinated vertically on half-strength Murashige and Skoog (MS) medium. All plant were grown under continuous white light (60 µmol/m^2^/s) at 24°C and 45% relative humidity.

### Immunolabeling and chemical staining of cell wall components

Immunolabeling of AGPs and pectins was performed essentially as described by (Berry et al., 2016). Protonemal tissues from 7-day-old *P. patens* cultures was collected and fixed in fixation buffer (50mM PIPES, 2.5 mM MgSO_4_, 5 mM EGTA, 4% paraformaldehyde) for 20 min. Samples were washed three times in phosphate-buffered saline (PBS; 137mM NaCl, 2.7mM KCl, 10mM Na_2_HPO_4_, 1.4mM KH_2_PO_4_) and mounted on poly-L-lysine-coated slides (Sigma-Aldrich, USA). Samples were blocked in 100 μL of 5% (w/v) skim milk in PBS for 10 min and incubated for 1.5 h at room temperature with primary antibodies diluted 1:5 in blocking solution: LM2 (AGP epitope; Kerafast, USA) or LM19 (unesterified homogalacturonan; Kerafast, USA). After three PBS washes, tissues were incubated for 1.5 h with Alexa Fluor 647-conjugated goat anti-rat IgG secondary antibody (1:100, Abcam, UK). Imaging was performed using a Zeiss LSM880 confocal microscope (Zeiss, Jena, Germany) equipped with a C-Apochromat 40×/1.2W Korr FCS M27 objective. Alexa Fluor 647 was excited at 633 nm, and emission was collected between 645–685 nm bandwidth.

For cellulose staining, 7-day-old protonemata were incubated in 0.1 mg/mL Calcofluor White (Sigma-Aldrich, USA) for 2 min prior to imaging. Calcofluor White was excited at 405 nm, and emission was collected within a 410–461 nm.

### Chemical inhibitors

Seven-day-old protonemata were preincubated with 2 μM FM4-64 (Thermo Fisher Scientific, USA) for 5 min. Brefeldin A (BFA) or Wortmannin (Wmn) were then added to final concentrations of 50 μM and 33 μM, respectively, followed by incubation of 1h. All samples were rinsed with water before imaging. For cellulose synthase inhibition assays, 6-day-old protonemal tissues were transferred to cellophane-overlain BCDAT agar medium supplemented with 0.02% v/v ethanol (mock control), 20 μM isoxaben (ISX), or 5 μM 2’, 6’-dichlorobenzonitrile (DCB), and grown for 10 days before subsequent phenotypic analysis.

## Imaging

### Confocal microscopy

For subcellular localization in Arabidopsis, 5-day-old AtEXP7_pro_:SB-mNG seedlings were stained with 5 μg/mL propidium iodide for 30 s and imaged using excitation at 488 nm (mNG) and 561 nm (propidium iodide). To visualize intracellular localization of SB-mNG or SB^P55,92,94A^-mNG, 7-day-old protonemata were stained with either 2 μM FM4-64 for 5 min or 2 μM MDY-64 for 10 min prior to inhibitor treatments and imaging. Samples were imaged using a Zeiss LSM880 confocal microscope equipped with a C-Apochromat 40×/1.2W Korr FCS M27 objective. Laser lines at 458 nm, 514 nm, and 633 nm were used to excite MDY-64, mNG, and chlorophyll, respectively. Emission was collected within 470–500 nm (MDY-64), 520–540 nm (mNG), and 660–690 nm (chlorophyll) using the Ch1 PMT, Ch2 GaAsP detector, and Ch3 PMT.

### Transmission electron microscopy (TEM)

Seven-day-old protonemata were fixed in 2.5 % v/v glutaraldehyde and 4 % w/v paraformaldehyde in 0.05 M sodium cacodylate buffer (pH 7.0) using microwave-assissted fixation (Pelco BioWave Pro+, TED PELLA Inc., USA). Samples were post-fixed in 1% (w/v) OsO₄, dehydrated through an ethanol series and propylene oxide, infiltrated with Spurr’s resin, and sectioned using a Leica EM UC6 ultramicrotome (Leica Microsystems, Germany). Ultrathin sections (70–90 nm) were stained with 5% w/v uranyl acetate and 0.4% w/v lead citrate and examined using a FEI Tecnai G2 Spirit Twin transmission electron microscope (Thermo Fisher Scientific, USA) operated at 80 kV.

### Brillouin microscopy

The detail of the instrumentation and data analysis can be found in our recent report(Alonso Baez et al., 2026). In brief, a Brillouin spectrometer with a two-stage VIPA configuration and a Lyott stop was built as an add-on module to a commercial Leica SP8 confocal microscope. Excitation was performed via a 532 nm laser (Cobolt). Light was collected in a backscattering configuration with an 40x/1.1 NA objective, and the spectra was recorded using a CMOS ORCA Quest camera (Hamamatsu). The laser power on the sample was 15mW and each spectrum was acquired by integrating 3 frames at 100ms. Samples were inspected before and after imaging using brightfield illumination and appeared unaffected and healthy. During spectra acquisition the microscope stage position was controlled by the HCImage software (Hamamatsu).

Data analysis was performed using a custom-developed MATLAB script to fit the spectra using a Lorentzian function (script available in (Alonso Baez et al., 2026)). Water and methanol were used as reference samples for the calculation of the frequency shift and linewidth.

### TIRF microscopy

To examine plasma membrane-localized SB-mNG or SBP55,92,94A-mNG, 7-day-old protonemata were imaged using a Zeiss LSM 980 inverted confocal plus super-resolution microscope (LSM 980 + ELYRA) with an α Plan-Apochromat 100×/1.46 Oil DIC M27 objective. Samples were excited with a 488 nm laser at a 60° angle under total internal reflection fluorescence (TIRF) mode, and emission was collected between 495–550 nm using a Hamamatsu camera (Hamamatsu Photonics, Japan).

### Time-lapse imaging

Six-day-old protonemata were immobilized in 35-mm glass-bottom dishes (ibidi GmbH, Germany) by overlaying with 0.6% (w/v) BCDAT agar and maintained overnight under standard growth conditions. Time-lapse imaging of growing apical cells was performed using a Leica THUNDER imaging system with images acquired at 30-min intervals. mNeonGreen was excited at 470 nm, and emission was collected using a 514 nm filter. Images were processed using Instant Computational Clearing in Leica Application Suite X to reduce out-of-focus blur.

### Image analysis

For *P. patens* growth analysis, 6-day-old protonemal tissues were transferred to inhibitor-containing or control media. Colony images were captured on day 1 and day 10 after treatment. Colony area was quantified using the Wand tool in ImageJ (v1.54p).

Bright-field images of 5-day-old Arabidopsis seedlings were acquired using a Leica M165 FC fluorescence stereomicroscope (Leica Microsystems, Germany). Root hair lengths were measured for the first ten consecutive root hairs using the “straight” selection tool in ImageJ software (v1.54p). Cuticular layer and cell wall thickness were quantified from TEM micrographs using eight evenly spaced measurements across the cuticular layer/cell wall per image; the average was treated as single biological data point.

Pearson’s correlation coefficients were calculated with ImageJ (v1.54p, NIH, USA) using the BIOP JACoP plugin (https://imagej.net/plugins/jacop).

### Immunoblotting

Total proteins were extracted from protonemata using 2 volumes of urea extraction buffer (4M Urea, 100mM DTT, 1% v/v Triton X-100) and separated by 7% or 10% SDS-PAGE. Immunoblotting was performed using rabbit polyclonal anti-mNG (Agrisera AS214525, 1:1,000), mouse monoclonal anti- 6x His (Yao-Hong Biotechnology YH80003, 1:1,000), followed by HRP-conjugated secondary anti-rabbit (Sigma-Aldrich A0545, 1:20,000) or anti-mouse (Agrisera AS09-627, 1:20,000).

### Microsomal membrane fractionation

Approximately 200 mg of 7-day-old protonemal tissue was ground in liquid nitrogen and homogenized in ice-cold buffer (50 mM HEPES-KOH pH 7.5, 250 mM sucrose, 5% glycerol, 10 mM EDTA, 0.5% polyvinylpyrrolidone, 50 mM sodium pyrophosphate, 1 mM sodium molybdate, 25 mM sodium fluoride, supplemented with 3 mM DTT, 1 mM PMSF, and 10 μM protease inhibitor cocktail). The homogenate was centrifuged at 8,000 g for 10 min. The supernatant (S_8_) was further centrifuged at 100,000 g for 30 min to obtain a soluble fraction (S_100_) and microsomal pellet (P_100_), which was washed once and resuspended in homogenization buffer. Fractions were analyzed by SDS-PAGE and immunoblotting using the following antibodies: anti-mNeonGreen (Agrisera AS214525, 1:2,000), anti-actin (Agrisera AS163141, 1:1,000), and anti-CNX1/2 (Agrisera AS12-2365, 1:2,500). HRP-conjugated secondary antibodies were used at 1:10,000.

### Molecular cloning

*Zm*UBQ:SB-mNG, *Zm*UBQ:SB^P55,92, 94A^-mNG, and AtEXP_pro_:SB-mNG constructs were generated by Golden Gate assembly (Marillonnet and Grutzner, 2020). An internal *Bsa*I site in the SB coding sequence was first domesticated by two fragments amplification of SB using primers #542/#543 and #544/#545, which were assembled into the Level-0 vector pAGM9121 to generate pOKT020003. The assembly reaction contained 1x T4 ligase buffer, 1x Buffer G, 5U *Bpi*I and 1.5U T4 DNA ligase. To ensure correct reading frame for downstream mocular cloning, pOKT020003 was further mutagenized using primers #579/#580 to produce pOKT020004. Sequence-verified pOKT020004 was then used as the SB Level-0 entry clone for Level-1 assembly using the same reaction buffer supplmented with 10U *Bsa*I. All Golden Gate reactions were cycled 40 times (3 min at 37°C, 4 min at 16°C), followed by 5 min incubations at 50°C and a final 5 min at 80°C. To generate hypoglycosylated SB variants, point mutations were introduced sequentially in pOKT020004, yielding pOKT020009 (SB^P55A^, primers #753/#754), pOKT020010 (SB^P55,92A^, primers #755/#756), and pOKT020011 (SB^P55,92, 94A^, primers #757/#758). These Level-0 constructs were subsequently used for Level-1 expression vector assembly. The 35S:SB-YFP construct was generated by LR recombination of pENTR1A:SB into pH35GY. All primers used in this study are listed in Supplementary Table 1.

## Acknowledgement

We thank Dr. Micheal Prigge (University of California San Diego, USA) for providing Golden Gate cloning parts. We are grateful to the IPMB Cell Biology Core Lab (Dr. Wann-Neng Jane), the G-Tec DNA Analysis Division (Ms. Me-Jane Fang), and the ABRC Advanced Optical Microscope Core Facility (Ms. Shu-Chen, Shen) for expert technical support.

This work was supported by the Academia Sinica Career Development Award Grant (AS-CDA-113-L04) and an IPMB intramural grant to O.-K. T. Additional support was provided by European Research Council (ERC HYDROSENSING, Project No. 101118769, to T.H.), The Research Council of Norway (WALLINTEGRITY Project no. 315325, to T.H.), NTNU Faculty of Natural Sciences (Brillouin microscope building grant to T.H.; Career Development Grant to L.A.B.).

## Supplementary figure legends

**Figure S1. SB overexpression inhibits colony growth (related to Figure 1)**

(A) Immunoblot analysis of mNG-immunoprecipitated proteins from independent SB-mNG transgenic lines.

(B) Immunoblot analysis of mNG-immunoprecipitated proteins following digestion with glycosidases PNGasF or EndoH.

(C) Images of four-week old WT and SB-mNG (lines #1 and #17) colonies grown on BCDAT or BCD medium. Scale bar, 5 mm.

(D) Images of seven-day-old colonies of WT, *sb sbl* and independent SB-mNG/*sb sbl* lines. Scale bar, 2 mm.

(E) Quantification of colony size after 7 days of growth on AT medium for WT (white circles), *sb sbl* (grey circles), and SB-mNG/sb sbl (grey circles with green boarder).

**Figure S2. Additional TIRF images of SB-mNG and SB^P55,92,94A^-mNG (related to Figure 2)**

(A, B) TIRF microscopy images of protonemal filanents expressing from SB-mNG (A) and SB^P55,92,94A^-mNG (B). Scale bar, 10 µm.

**Figure S3. SB-mNG does not undergo endosomal recycling (related to Figure 3)**

(A, B) Confocal images of apical protonemata expressing SB-mNG (A) or SB^P55,92,94A^-mNG (B) after treatment with 50 µM BFA. Images are single optical sections. Yellow arrowheads indicate BFA compartments. Blue, green and magenta denote chloroplast autofluorescence, mNG and FM4-64, respectively. Scale bars, 5 µm.

**Figure S4. Cell wall ultrastructure and mechanical properties in SB alleles (related to Figure 4)**

(A) Representative TEM micrographs of WT (top) and *sb sbl* with an intact cuticular layer (CL; middle) or lacking CL (bottom). Numbers in parentheses indicate the fraction of micrographs displaying the indicated morphology. CP, cuticle proper; CL, cuticular later; CW, cell wall; PM, plasma membrane. Scale bar, 200 nm.

(B) Scatter dot plot of cuticular layer thickness in WT (white circles), *sb sbl* (grey circles and triangles), SB-mNG (green circles), and SB^P55,92,94A^-mNG (mangenta circles). Triangles indicate measurements from *sb sbl* micrographs retaining a CL. Different letters indicate statistically significant differences (one-way ANOVA with Tukey’s HSD test, *α*=0.05).

(C) Confocal images of protonemata immunolabelled with the LM19 antibody. Images are z-stack projections (sum intensity). Asterisks indicate autofluoresence from collapsed protoplasts. Scale bar, 10 µm.

(D) Schematic representation of Brillouin shift measurements taken from the apical cell outline (blue) and central region (orange).

(E, F) Representative heatmaps of Brillouin linewidths in apical tip cells (F) and corresponding quantification (E). Measurements were obtained from WT (white), *sb sbl* (grey), and SB-mNG (green) in the tip cell outline (circles) and central regions (diamond). Statistical analysis was performed using ANOVA with Šidák’s multiple-comparison test (*α*=0.05). n.s., not significant.

**Figure S5. Isoxaben (ISX) treatment responses of SB alleles (related to Figure 5)**

(A) Representative 10-day old colonies of WT, *sb sbl*, SB-mNG, and SB^P55,92,94A^-mNG grown under mock conditions (left) or in the presence of 20 µM ISX (right). Scale bar, 2 mm.

(B) Scatter plot of colony size under mock conditions (circles) or 20 µM ISX treatment (squares). Different letters indicate statistically significant differences between groups (one-way ANOVA with Tukey’s HSD, *α*=0.05).

**Figure S6. Cellular geometry of SB alleles (related to Figure 6)**

(A, B) Scatter plot of cell length (A) and cell width (B) in WT (white circles), *sb sbl* (grey circles), SB-mNG (green circles), and SB^P55,92,94A^-mNG (mangenta circles). Different letters indicate statistically significant differences (one-way ANOVA with Tukey’s multiple-comparison test, *α*=0.05).

(C) Confocal image of an actively dividing gametophore leaf. Scale bar, 20 µm.

**Figure S7. SB overexpression effects in Arabidopsis (related to Figure 7)**

(A) Bright-field images of 35-day-old Arabidopsis plants from Col-0, *35S:YFP* and *35S:SB-YFP* lines. Scale bar, 5 cm.

(B) Scatter plot of inflorescence length in Col-0 (white circles), *35S:YFP* (yellow triangles) and *35S:SB-YFP* (yellow circles). Mean values are indicated by dotted lines, error bars represent standard deviation. Letters denote statistically sgnificant differences (one-way ANOVA with Tukey’s HSD test, *α*=0.05).

(C) Immunoblots detecting YFP and Calnexin (CNX) in total (T), microsomal (P), and soluble (S) protein fractions from Col-0 (white circles), *35S:YFP* (yellow triangles), and *35S:SB-YFP* (yellow circles).

## References

Alonso Baez L, Bjørkøy A, Saffioti F, Morghen S, Amanda D, Tichá M, Besten M, Ivanova A, Sprakel J, Stokke BT, Hamann T (2026) The mechanical properties of Arabidopsis thaliana roots adapt dynamically during development and to stress.

Anderson JR, Barnes WS, Bedinger P (2002) 2,6-dichlorobenzonitrile, a cellulose biosynthesis inhibitor, affects morphology and structural integrity of petunia and lily pollen tubes. Journal of Plant Physiology 159: 61–67

Bacete L, Schulz J, Engelsdorf T, Bartosova Z, Vaahtera L, Yan G, Gerhold JM, Ticha T, Ovstebo C, Gigli-Bisceglia N, Johannessen-Starheim S, Margueritat J, Kollist H, Dehoux T, McAdam SAM, Hamann T (2022) THESEUS1 modulates cell wall stiffness and abscisic acid production in Arabidopsis thaliana. Proc Natl Acad Sci U S A 119: e2119258119

Berry EA, Tran ML, Dimos CS, Budziszek MJ, Jr., Scavuzzo-Duggan TR, Roberts AW (2016) Immuno and Affinity Cytochemical Analysis of Cell Wall Composition in the Moss Physcomitrella patens. Front Plant Sci 7: 248

Bibeau JP, Galotto G, Wu M, Tuzel E, Vidali L (2021) Quantitative cell biology of tip growth in moss. Plant Mol Biol 107: 227–244

Borassi C, Gloazzo Dorosz J, Ricardi MM, Carignani Sardoy M, Pol Fachin L, Marzol E, Mangano S, Rodriguez Garcia DR, Martinez Pacheco J, Rondon Guerrero YDC, Velasquez SM, Villavicencio B, Ciancia M, Seifert G, Verli H, Estevez JM (2020) A cell surface arabinogalactan-peptide influences root hair cell fate. New Phytol 227: 732–743

Borner GH, Sherrier DJ, Weimar T, Michaelson LV, Hawkins ND, Macaskill A, Napier JA, Beale MH, Lilley KS, Dupree P (2005) Analysis of detergent-resistant membranes in Arabidopsis. Evidence for plasma membrane lipid rafts. Plant Physiol 137: 104–116

Bove J, Vaillancourt B, Kroeger J, Hepler PK, Wiseman PW, Geitmann A (2008) Magnitude and direction of vesicle dynamics in growing pollen tubes using spatiotemporal image correlation spectroscopy and fluorescence recovery after photobleaching. Plant Physiol 147: 1646–1658

Cho HT, Cosgrove DJ (2002) Regulation of root hair initiation and expansin gene expression in Arabidopsis. Plant Cell 14: 3237–3253

Cove DJ, Perroud PF, Charron AJ, McDaniel SF, Khandelwal A, Quatrano RS (2009) Transformation of the moss Physcomitrella patens using direct DNA uptake by protoplasts. Cold Spring Harb Protoc 2009: pdb prot5143

Dehors J, Mareck A, Kiefer-Meyer MC, Menu-Bouaouiche L, Lehner A, Mollet JC (2019) Evolution of Cell Wall Polymers in Tip-Growing Land Plant Gametophytes: Composition, Distribution, Functional Aspects and Their Remodeling. Front Plant Sci 10: 441

Favery B, Ryan E, Foreman J, Linstead P, Boudonck K, Steer M, Shaw P, Dolan L (2001) KOJAK encodes a cellulose synthase-like protein required for root hair cell morphogenesis in Arabidopsis. Genes Dev 15: 79–89

Feng X, Zheng J, Irisarri I, Yu H, Zheng B, Ali Z, de Vries S, Keller J, Furst-Jansen JMR, Dadras A, Zegers JMS, Rieseberg TP, Dhabalia Ashok A, Darienko T, Bierenbroodspot MJ, Gramzow L, Petroll R, Haas FB, Fernandez-Pozo N, Nousias O, Li T, Fitzek E, Grayburn WS, Rittmeier N, Permann C, Rumpler F, Archibald JM, Theissen G, Mower JP, Lorenz M, Buschmann H, von Schwartzenberg K, Boston L, Hayes RD, Daum C, Barry K, Grigoriev IV, Wang X, Li FW, Rensing SA, Ben Ari J, Keren N, Mosquna A, Holzinger A, Delaux PM, Zhang C, Huang J, Mutwil M, de Vries J, Yin Y (2024) Genomes of multicellular algal sisters to land plants illuminate signaling network evolution. Nat Genet

Furt F, Lemoi K, Tuzel E, Vidali L (2012) Quantitative analysis of organelle distribution and dynamics in Physcomitrella patens protonemal cells. BMC Plant Biol 12: 70

Geldner N, Anders N, Wolters H, Keicher J, Kornberger W, Muller P, Delbarre A, Ueda T, Nakano A, Jurgens G (2003) The Arabidopsis GNOM ARF-GEF mediates endosomal recycling, auxin transport, and auxin-dependent plant growth. Cell 112: 219–230

Grefen C, Donald N, Hashimoto K, Kudla J, Schumacher K, Blatt MR (2010) A ubiquitin-10 promoter-based vector set for fluorescent protein tagging facilitates temporal stability and native protein distribution in transient and stable expression studies. Plant J 64: 355–365

Gu Y, Fu Y, Dowd P, Li S, Vernoud V, Gilroy S, Yang Z (2005) A Rho family GTPase controls actin dynamics and tip growth via two counteracting downstream pathways in pollen tubes. J Cell Biol 169: 127–138

Hao H, Chen T, Fan L, Li R, Wang X (2013) 2, 6-Dichlorobenzonitrile causes multiple effects on pollen tube growth beyond altering cellulose synthesis in Pinus bungeana Zucc. PLoS One 8: e76660

Hijazi M, Velasquez SM, Jamet E, Estevez JM, Albenne C (2014) An update on post-translational modifications of hydroxyproline-rich glycoproteins: toward a model highlighting their contribution to plant cell wall architecture. Front Plant Sci 5: 395

Hromadova D, Soukup A, Tylova E (2021) Arabinogalactan Proteins in Plant Roots - An Update on Possible Functions. Front Plant Sci 12: 674010

Hu G, Katuwawala A, Wang K, Wu Z, Ghadermarzi S, Gao J, Kurgan L (2021) flDPnn: Accurate intrinsic disorder prediction with putative propensities of disorder functions. Nat Commun 12: 4438

Kim SJ, Brandizzi F (2016) The plant secretory pathway for the trafficking of cell wall polysaccharides and glycoproteins. Glycobiology 26: 940–949

Lamport DT, Varnai P, Seal CE (2014) Back to the future with the AGP-Ca2+ flux capacitor. Ann Bot 114: 1069–1085

Lamport DTA, Varnai P (2013) Periplasmic arabinogalactan glycoproteins act as a calcium capacitor that regulates plant growth and development. New Phytol 197: 58–64

Lee KJ, Sakata Y, Mau SL, Pettolino F, Bacic A, Quatrano RS, Knight CD, Knox JP (2005) Arabinogalactan proteins are required for apical cell extension in the moss Physcomitrella patens. Plant Cell 17: 3051–3065

Lee YJ, Szumlanski A, Nielsen E, Yang Z (2008) Rho-GTPase-dependent filamentous actin dynamics coordinate vesicle targeting and exocytosis during tip growth. J Cell Biol 181: 1155–1168

Li J, Fan L, Yang T, Zhang P, Ruan H, Li Y, Wang T, Zhang Y, Zhang F, Ren H (2024) AtFH5 recruits and transports the arabinogalactan protein AGP23 to maintain the tip growth of pollen tube. Proc Natl Acad Sci U S A 121: e2410607121

Lin S, Miao Y, Huang H, Zhang Y, Huang L, Cao J (2022) Arabinogalactan Proteins: Focus on the Role in Cellulose Synthesis and Deposition during Plant Cell Wall Biogenesis. Int J Mol Sci 23

Lopez-Hernandez F, Tryfona T, Rizza A, Yu XL, Harris MOB, Webb AAR, Kotake T, Dupree P (2020) Calcium Binding by Arabinogalactan Polysaccharides Is Important for Normal Plant Development. Plant Cell 32: 3346–3369

Luo N, Yan A, Liu G, Guo J, Rong D, Kanaoka MM, Xiao Z, Xu G, Higashiyama T, Cui X, Yang Z (2017) Exocytosis-coordinated mechanisms for tip growth underlie pollen tube growth guidance. Nat Commun 8: 1687

Ma Y, Johnson K (2023) Arabinogalactan proteins - Multifunctional glycoproteins of the plant cell wall. Cell Surf 9: 100102

MacAlister CA, Ortiz-Ramirez C, Becker JD, Feijo JA, Lippman ZB (2016) Hydroxyproline O-arabinosyltransferase mutants oppositely alter tip growth in Arabidopsis thaliana and Physcomitrella patens. Plant J 85: 193–208

Malivert A, Erguvan O, Chevallier A, Dehem A, Friaud R, Liu M, Martin M, Peyraud T, Hamant O, Verger S (2021) FERONIA and microtubules independently contribute to mechanical integrity in the Arabidopsis shoot. PLoS Biol 19: e3001454

Marillonnet S, Grutzner R (2020) Synthetic DNA Assembly Using Golden Gate Cloning and the Hierarchical Modular Cloning Pipeline. Curr Protoc Mol Biol 130: e115

Molendijk AJ, Bischoff F, Rajendrakumar CS, Friml J, Braun M, Gilroy S, Palme K (2001) Arabidopsis thaliana Rop GTPases are localized to tips of root hairs and control polar growth. EMBO J 20: 2779–2788

Munch D, Teh OK, Malinovsky FG, Liu Q, Vetukuri RR, El Kasmi F, Brodersen P, Hara-Nishimura I, Dangl JL, Petersen M, Mundy J, Hofius D (2015) Retromer contributes to immunity-associated cell death in Arabidopsis. Plant Cell 27: 463–479

Park HO, Bi EF (2007) Central roles of small GTPases in the development of cell polarity in yeast and beyond. Microbiology and Molecular Biology Reviews 71: 48–96

Parsons HT, Stevens TJ, McFarlane HE, Vidal-Melgosa S, Griss J, Lawrence N, Butler R, Sousa MML, Salemi M, Willats WGT, Petzold CJ, Heazlewood JL, Lilley KS (2019) Separating Golgi Proteins from Cis to Trans Reveals Underlying Properties of Cisternal Localization. Plant Cell 31: 2010–2034

Petersen BL, MacAlister CA, Ulvskov P (2021) Plant Protein O-Arabinosylation. Front Plant Sci 12: 645219

Radin I, Richardson RA, Coomey JH, Weiner ER, Bascom CS, Li T, Bezanilla M, Haswell ES (2021) Plant PIEZO homologs modulate vacuole morphology during tip growth. Science 373: 586–590

Roberts EM, Yuan K, Chaves AM, Pierce ET, Cresswell R, Dupree R, Yu X, Blanton RL, Wu SZ, Bezanilla M, Dupree P, Haigler CH, Roberts AW (2024) An alternate route for cellulose microfibril biosynthesis in plants. Sci Adv 10: eadr5188

Samalova M, Melnikava A, Elsayad K, Peaucelle A, Gahurova E, Gumulec J, Spyroglou I, Zemlyanskaya EV, Ubogoeva EV, Balkova D, Demko M, Blavet N, Alexiou P, Benes V, Mouille G, Hejatko J (2023) Hormone-regulated expansins: Expression, localization, and cell wall biomechanics in Arabidopsis root growth. Plant Physiol 194: 209–228

Shimizu M, Igasaki T, Yamada M, Yuasa K, Hasegawa J, Kato T, Tsukagoshi H, Nakamura K, Fukuda H, Matsuoka K (2005) Experimental determination of proline hydroxylation and hydroxyproline arabinogalactosylation motifs in secretory proteins. Plant J 42: 877–889

Showalter AM, Keppler B, Lichtenberg J, Gu D, Welch LR (2010) A bioinformatics approach to the identification, classification, and analysis of hydroxyproline-rich glycoproteins. Plant Physiol 153: 485–513

Silva J, Ferraz R, Dupree P, Showalter AM, Coimbra S (2020) Three Decades of Advances in Arabinogalactan-Protein Biosynthesis. Front Plant Sci 11: 610377

Storti M, Costa A, Golin S, Zottini M, Morosinotto T, Alboresi A (2018) Systemic Calcium Wave Propagation in Physcomitrella patens. Plant Cell Physiol 59: 1377–1384

Tan L, Eberhard S, Pattathil S, Warder C, Glushka J, Yuan C, Hao Z, Zhu X, Avci U, Miller JS, Baldwin D, Pham C, Orlando R, Darvill A, Hahn MG, Kieliszewski MJ, Mohnen D (2013) An Arabidopsis cell wall proteoglycan consists of pectin and arabinoxylan covalently linked to an arabinogalactan protein. Plant Cell 25: 270–287

Tan L, Showalter AM, Egelund J, Hernandez-Sanchez A, Doblin MS, Bacic A (2012) Arabinogalactan-proteins and the research challenges for these enigmatic plant cell surface proteoglycans. Frontiers in Plant Science 3

Teh OK, Singh P, Ren J, Huang LT, Ariyarathne M, Salamon BP, Wang Y, Kotake T, Fujita T (2022) Surface-localized glycoproteins act through class C ARFs to fine-tune gametophore initiation in Physcomitrium patens. Development 149: dev200370

Vaahtera L, Schulz J, Hamann T (2019) Cell wall integrity maintenance during plant development and interaction with the environment. Nat Plants 5: 924–932

Velasquez SM, Ricardi MM, Dorosz JG, Fernandez PV, Nadra AD, Pol-Fachin L, Egelund J, Gille S, Harholt J, Ciancia M, Verli H, Pauly M, Bacic A, Olsen CE, Ulvskov P, Petersen BL, Somerville C, Iusem ND, Estevez JM (2011) O-Glycosylated Cell Wall Proteins Are Essential in Root Hair Growth. Science 332: 1401–1403

Wang J, Cai Y, Miao Y, Lam SK, Jiang L (2009) Wortmannin induces homotypic fusion of plant prevacuolar compartments. J Exp Bot 60: 3075–3083

Wang L, Cheng M, Yang Q, Li J, Wang X, Zhou Q, Nagawa S, Xia B, Xu T, Huang R, He J, Li C, Fu Y, Liu Y, Bao J, Wei H, Li H, Tan L, Gu Z, Xia A, Huang X, Yang Z, Deng XW (2019) Arabinogalactan protein-rare earth element complexes activate plant endocytosis. Proc Natl Acad Sci U S A 116: 14349–14357

Wu SZ, Chaves AM, Li R, Roberts AW, Bezanilla M (2023) Cellulose synthase-like D movement in the plasma membrane requires enzymatic activity. J Cell Biol 222

Xue H, Veit C, Abas L, Tryfona T, Maresch D, Ricardi MM, Estevez JM, Strasser R, Seifert GJ (2017) Arabidopsis thaliana FLA4 functions as a glycan-stabilized soluble factor via its carboxy-proximal Fasciclin 1 domain. Plant J 91: 613–630

Yeats TH, Rose JK (2013) The formation and function of plant cuticles. Plant Physiol 163: 5–20

Yi P, Goshima G (2020) Rho of Plants GTPases and Cytoskeletal Elements Control Nuclear Positioning and Asymmetric Cell Division during Physcomitrella patens Branching. Curr Biol 30: 2860–2868 e2863

Yuasa K, Toyooka K, Fukuda H, Matsuoka K (2005) Membrane-anchored prolyl hydroxylase with an export signal from the endoplasmic reticulum. Plant Journal 41: 81–94

Zhang N, Wright T, Wang X, Karki U, Savary BJ, Xu J (2019) Engineering ‘designer’ glycomodules for boosting recombinant protein secretion in tobacco hairy root culture and studying hydroxyproline-O-glycosylation process in plants. Plant Biotechnol J 17: 1130–1141

Zhou Z, Shi H, Chen B, Zhang R, Huang S, Fu Y (2015) Arabidopsis RIC1 Severs Actin Filaments at the Apex to Regulate Pollen Tube Growth. Plant Cell 27: 1140–1161

