## Supplementary data for "Intracellular trafficking of an Arabinogalactan protein SLEEPING BEAUTY influences cell wall integrity and apical tip growth in *Physcomitrium patens*"

### Figure S1 (related to Figure 1)

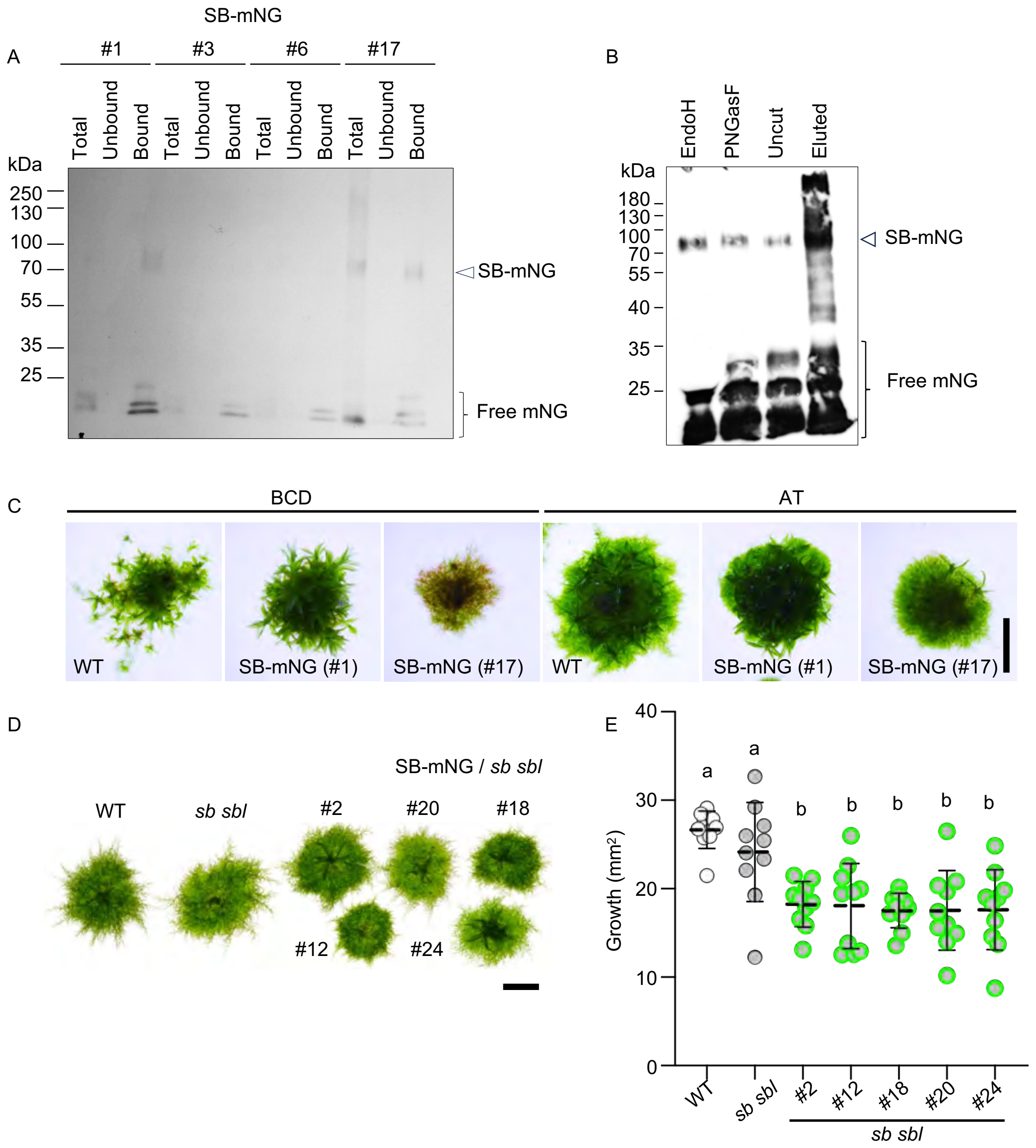

### Figure S2 (related to Figure 2)

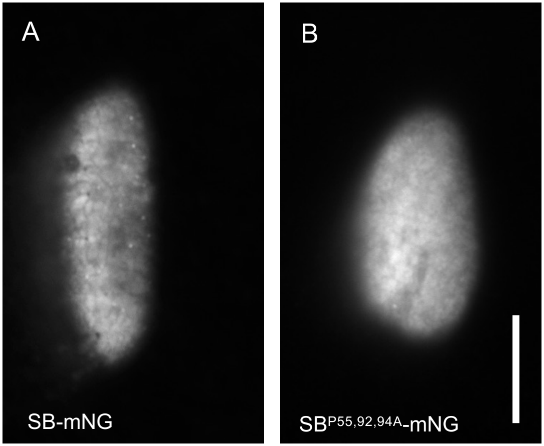

### Figure S3 (related to Figure 3)

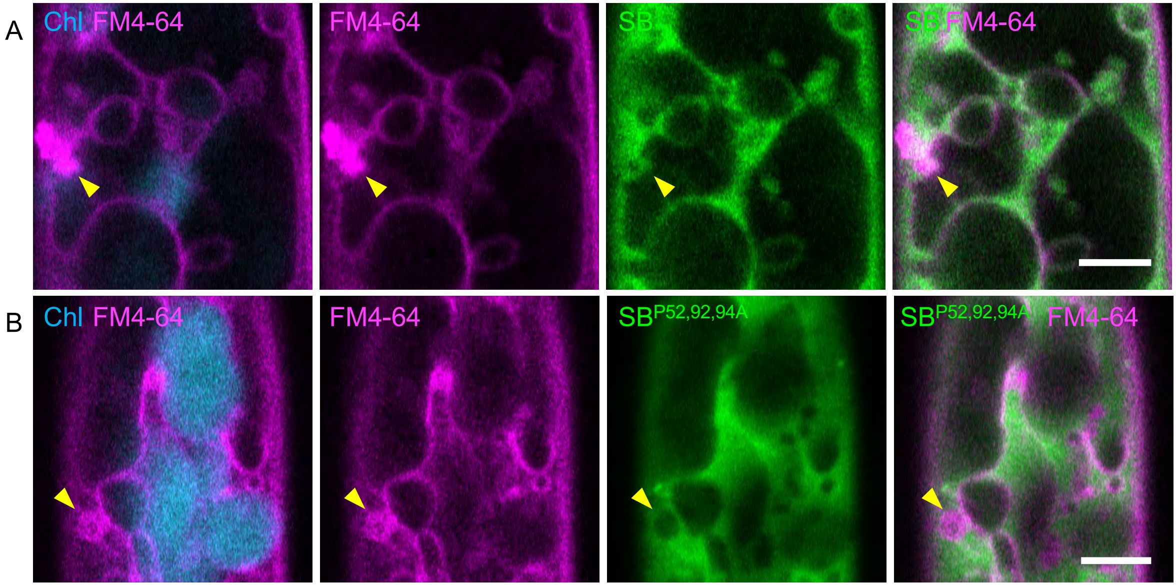

### Figure S4 (related to Figure 4)

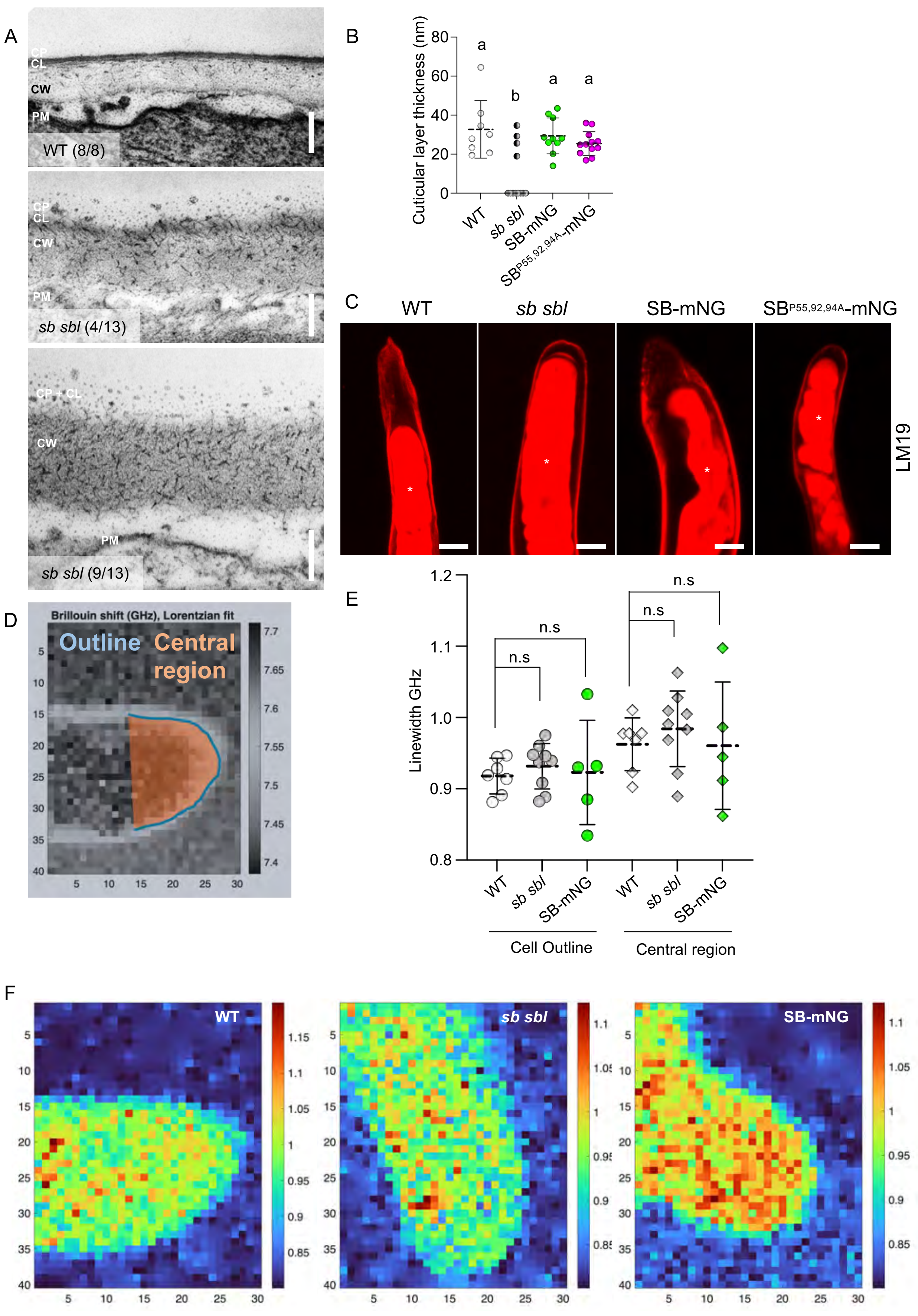

### Figure S5 (related to Figure 5)

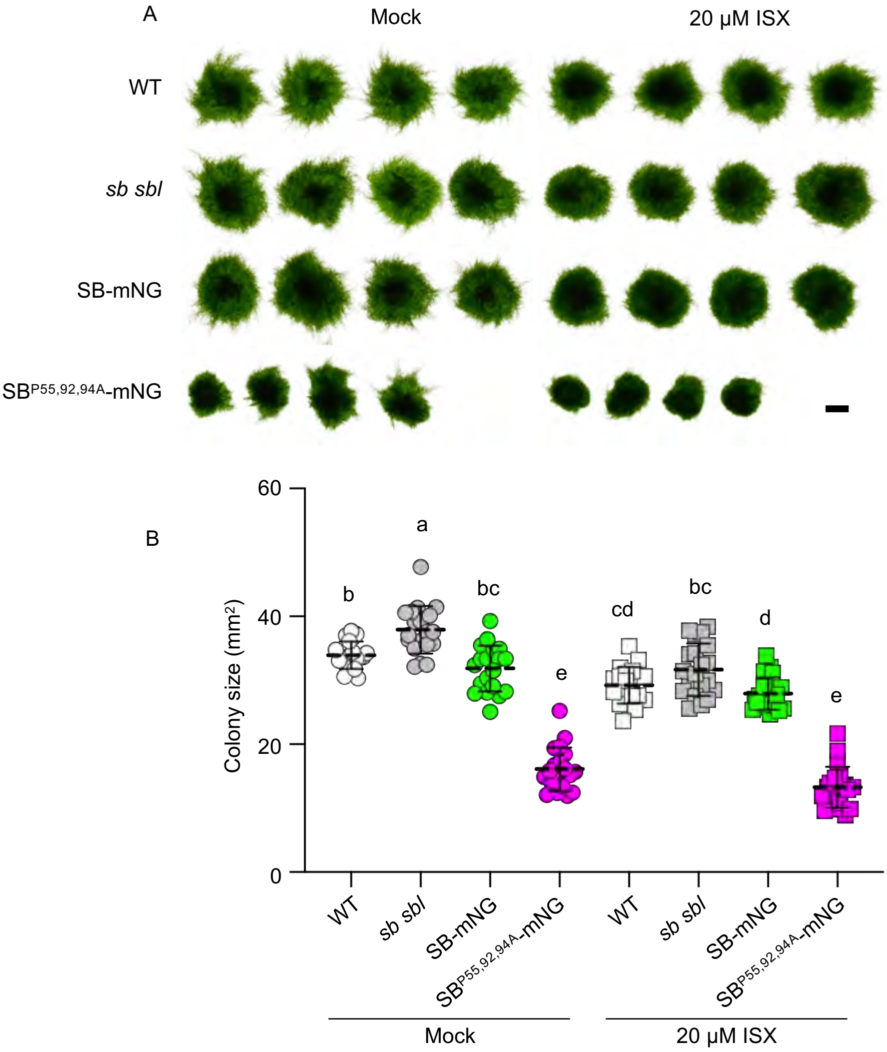

### Figure S6 (related to Figure 6)

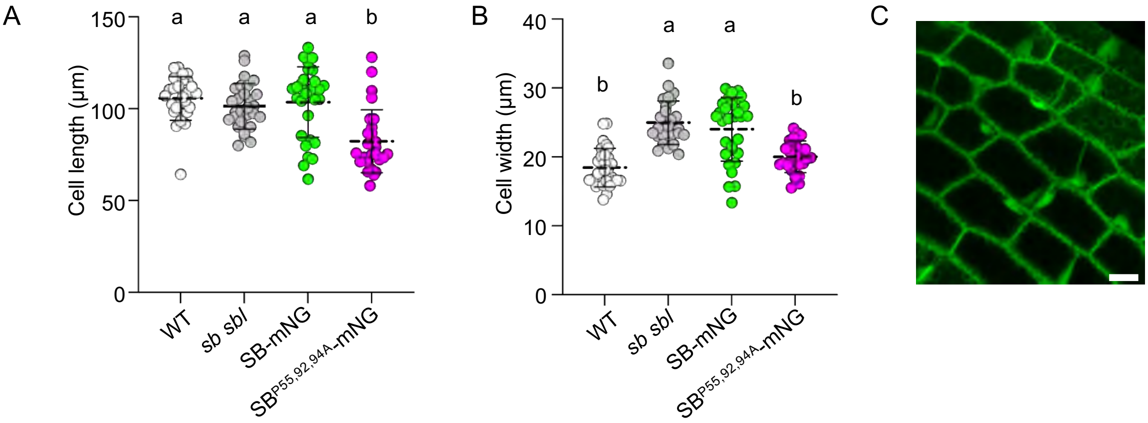

### Figure S7 (related to Figure 7)

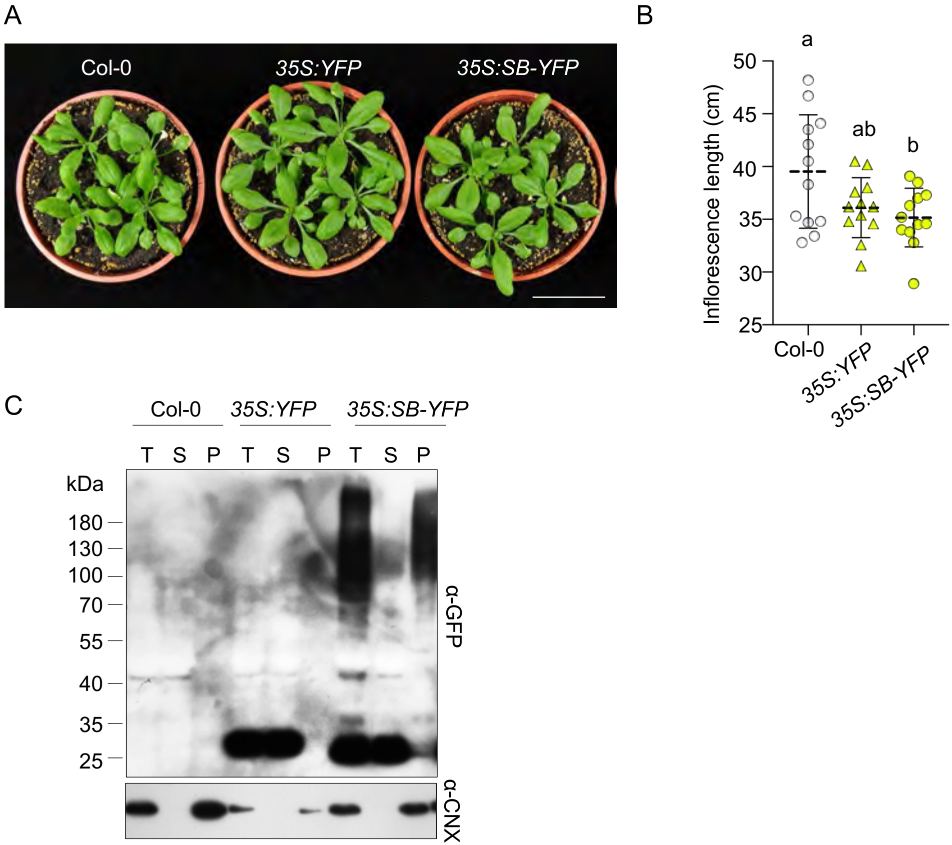

#### Supplementary Table 1

| Primer Number | Oligonucleotide sequence (5'-3') | Purpose |
|---|---|---|
| #542 | TT GAAGAC AT CTCA AATG ATGGCGGGTTCGTGGCGAGGTT | Domestication of SB |
| #543 | TT GAAGAC AT TCCC CGAATTTGCGGCAGAGGCAGAA | Domestication of SB |
| #579 | TTACCACTCTTCGCGAGACCACGAAGTGCC | Mutagenesis of pOKT020003 |
| #580 | TCTCGCGAAGAGTGGTAACGAGGCAAAGGC | Mutagenesis of pOKT020003 |
| #753 | GTGTT GCT GCCACCCCCCCTGTTTCTTC | Introduce P55A mutation |
| #754 | GTGGC AGC AACACCACCAGCCTGAAG | Introduce P55A mutation |
| #755 | CTGGTGCAGCTCCAGCTCCAGCTCCAGCTGC | Introduce P92A mutation |
| #756 | CTGGAGC TGC ACCAGCAGCCTTCGGAGG | Introduce P92A mutation |
| #757 | TGCAGCT GCA GCTCCAGCTCCAGCTGCCCC | Introduce P94A mutation |
| #758 | TGGAGC TGC AGCTGCACCAGCAGCCTTC | Introduce P94A mutation |
